# Phylogenomics recovers multiple origins of portable case-making in caddisflies (Insecta: Trichoptera), the world’s most common underwater architects

**DOI:** 10.1101/2023.12.21.572910

**Authors:** Paul B. Frandsen, Ralph W. Holzenthal, Marianne Espeland, Jesse Breinholt, Jessica A. Thomas, Sabrina Simon, Akito Y. Kawahara, David Plotkin, Scott Hotaling, Yiyuan Li, C. Riley Nelson, Oliver Niehuis, Christoph Mayer, Lars Podsiadlowski, Alex Donath, Bernhard Misof, Emily Moriarty Lemmon, Alan Lemmon, John C. Morse, Steffen Pauls, Xin Zhou

**Author notes:** correspondence: Paul B. Frandsen.

## Abstract

Caddisflies (Trichoptera) are among the most diverse groups of freshwater animals with more than 16,000 described species. They play an outsized role in freshwater ecology and environmental engineering in streams, rivers, and lakes. Because of this, they are frequently used as indicator organisms in biomonitoring programs. Despite their importance, key questions concerning the evolutionary history of caddisflies, such as the timing and origin of larval case-making, have been unanswered due to the lack of a well-resolved phylogenetic tree. To shed light on these questions in Trichoptera, we estimated a phylogenetic tree using a combination of transcriptomes and targeted enrichment data for 206 species, representing 48 of 52 extant families and 174 genera. We calibrated and dated the tree with a set of 33 carefully selected fossils. The first caddisflies originated in the Permian and the major suborders began to diversify in the Triassic. Ancestral state reconstruction and diversification analysis revealed that portable case-making evolved in three separate lineages and shifts in diversification occurred in concert with key evolutionary innovations other than case-making.

## Introduction

Freshwater is one of the world’s most limited and precious resources. It is inextricably linked to the survival and diversity of a multitude of terrestrial and freshwater species. To understand how to preserve this precious resource, decades of freshwater biomonitoring researchers have used aquatic macroinvertebrates to measure the health of streams. This is because macroinvertebrates are abundant, diverse, and differentially sensitive to environmental perturbation. Roughly 10% of the ~1 million described insect species on Earth spend at least one life stage in water (Hotaling et al. 2020) and have transitioned from terrestrial habitats to freshwater habitats on nearly 50 occasions (Dijkstra et al. 2014). Inhabiting rivers and lakes on six continents, these aquatic insects play critical roles in community ecology and ecosystem services (Morse et al. 2019). Five insect orders are almost exclusively aquatic (i.e., they require water for their entire larval development): Ephemeroptera (mayflies), Megaloptera (alderflies, dobsonflies, and fishflies), Odonata (dragonflies), Plecoptera (stoneflies), and Trichoptera (caddisflies). Of these, Trichoptera is the most species-rich, comprising more than 16,000 species (Morse 2009), making them the second most species-diverse radiation in all of the freshwater animals (Malm et al. 2013).

In addition to being incredibly diverse, caddisflies are perhaps the most common underwater architects on Earth. Most caddisfly larvae build composite underwater structures, which are often portable, out of a wide array of local materials, including rocks, sticks, and leaves (Wiggins 2005). The key adaptation underlying this behavior is silk production, which they share with their sister order, Lepidoptera (butterflies and moths). The sister order relationship between Lepidoptera and Trichoptera is among the most strongly supported within Insecta (Misof et al. 2014) and together they are known as Amphiesmenoptera. However, in contrast to the aquatic environments that caddisflies inhabit, the vast majority of Lepidoptera are terrestrial, with both orders using silk production in different ways. Caddisflies use silk to dramatically alter the physical environments of aquatic ecosystems, primarily at the interface between streambeds and the hyporheic zones beneath them. For instance, net-spinning caddisflies (family Hydropsychidae) frequently occur at densities of thousands per square meter (Miller 1984) and modify the stream environment in two ways; through nets constructed in the interstitial spaces of stream substrate to capture organic particles and by building retreats they live within (Statzner et al. 1999). Through these activities, net-spinning caddisflies can reduce vertical hydraulic conductivity of the streambed--a measure of permeability--by 55% (MacDonald et al. 2021). Ecosystem engineering by caddisflies likely extends beyond physical habitat modification to also include biogeochemical cycling. In experimental streams, the presence of caddisflies reduced in-stream organic matter by 50% (MacDonald et al. 2021). Despite their outsized importance in both freshwater ecosystem function and practical use as bioindicator organisms, questions remain concerning their evolutionary history, limiting our understanding of the evolutionary basis of these key functions.

Early systematic studies were based primarily on morphology, and were conducted before current analytical methods were available (Ross 1967; Weaver III 1984; Wiggins and Wichard 1989; Frania and Wiggins 1997). More recent studies based on a handful of molecular markers or mitochondrial genomes generated conflicting results (Malm et al. 2013; Thomas et al. 2020; Ge et al. 2023). Caddisflies are currently grouped into two major clades (suborders) reflecting the extended phenotypes of their larvae: Integripalpia and Annulipalpia. Integripalpia (Thomas et al. 2020) includes portable case-makers and free-living caddisflies, while Annulipalpia includes the so-called “fixed retreat makers”, whose larvae build fixed structures, usually in flowing waters, on rocks or other submerged substrates. The architectures created by species in these clades vary widely (Wiggins 2005). Understanding the timing and pattern of Trichoptera evolution is essential to interpreting the role of silk production and case-making in the diversification of the order. However, despite recent results that resolve many evolutionary relationships in caddisflies, incongruence among hypotheses still exists primarily with regard to the placement of five families. These are: the predatory, free-living caddisflies in the families Rhyacophilidae and Hydrobiosidae, the tortoise-case makers in the family Glossosomatidae, and two families of micro-caddisflies, Hydroptilidae and Ptilocolepidae. These families have been included in a group called “Spicipalpia” (Weaver III 1984; Wiggins and Wichard 1989), “closed cocoon-makers,” or simply “cocoon-makers” due to their construction of semi-permeable closed cocoons (Wiggins 2007). However, most Trichoptera researchers suspect that this group is not monophyletic, as a morphological assessment (Ivanov 1997) and all molecular estimates have recovered this grouping as para- or polyphyletic (Kjer et al. 2001, 2016; Malm et al. 2013;

Thomas et al. 2020; Ge et al. 2023). Because these families arose during the early evolution of the order and represent a diversity of silk-use and case-making strategies, understanding their correct placement in the evolutionary history of caddisflies is vital toward determining the origin of case-making and other behaviors of Trichoptera. Additionally, resolving the timing of the origin and diversification of caddisflies is essential for building a better understanding of the innovations that led to the evolutionary success of this critical group of freshwater insects.

To shed light on these questions, we generated a phylogenomic dataset aimed at estimating the evolutionary history of caddisflies. We combined *de novo* transcriptome sequencing, targeted exon capture sequencing, and publicly available genome assemblies to generate a data set for 206 caddisfly species representing 174 genera and 48 of 52 extant families. We resolve the timing and pattern of the evolutionary history of Trichoptera, including key nodes that have long been a systematic difficulty. In addition, we reconstruct ancestral states to determine the ancestral condition of larval retreat and case-making and pupal constructions. We then link our phylogenetic results to the evolution of the incredible array of underwater architectures constructed by caddisflies, thereby providing a key link between evolutionary history and phenotypic diversity that impacts the physical structure, hydrology, and ecology of freshwaters worldwide.

## Results

### Transcriptome results

Our final alignment in the transcriptome-only dataset consisted of 1,253,378 amino acids from 3,206 orthologous genes. The maximum likelihood (ML) tree based on the concatenated and partitioned supermatrix was largely congruent with the species tree generated in ASTRAL, with a few exceptions. Both trees recovered Annulipalpia and Integripalpia as reciprocally monophyletic. As in Thomas et al. (Thomas et al. 2020), the five cocoon-making families form a grade leading to the tube-case makers, Phryganides (Integripalpia, *s*.*l*.). The microcaddisflies, Ptilocolepidae and Hydroptilidae, form a monophyletic clade and are sister to the rest of Integripalpia. A clade containing the free-living (Rhyacophilidae and Hydrobiosidae) and tortoise-case making (Glossosomatidae) caddisflies is then sister to the tube case makers, subterorder Phryganides. While this clade has not been recovered in previous phylogenetic analyses, these families have been hypothesized to be closely related based on their pupal behavior, including the construction of a pupal dome, which protects a silken semi-permeable cocoon (Wiggins 2005). The most notable areas of incongruence between the ASTRAL species tree and the supermatrix-based ML tree are the relationships within this clade. In the supermatrix-based tree, the free-living caddisfly family Hydrobiosidae is recovered as sister to a clade containing the free-living family Rhyacophilidae and the tortoise-case making family Glossosomatidae. However, in the ASTRAL multispecies coalescent (MSC) tree, the two free-living families are recovered as a clade, sister to the tortoise-case makers. Both nodes are recovered with lower support than the other relationships in the tree. The bootstrap support in the transcriptome tree was 67, while the local posterior probability support in the MSC tree was 0.96. To further investigate this difference, we conducted four-cluster likelihood mapping with permutation tests on the transcriptome-only dataset (Fig. S1). Four-cluster likelihood mapping recovered a slight preference for (Hydrobiosidae + (Glossosomatidae + Rhyacophilidae)) over the alternative. However, the permutation tests revealed that (Hydrobiosidae + (Glossosomatidae + Rhyacophilidae)) was also the preferred arrangement in permutation test #1, revealing that this arrangement was driven by bias in the supermatrix-derived tree due to among-lineage heterogeneity and non-random distribution of missing data.

### Combined dataset

To boost the taxon sampling of the transcriptome data set, we sequenced an additional 158 individuals using targeted enrichment sequencing. After filtering, the final data set consisted of 339 orthologs representing 24,678 aligned amino acids. The deep splits in the maximum-likelihood tree based on the partitioned supermatrix were largely congruent with those recovered in the transcriptome-only tree (Fig. 1). However, contrary to the phylogeny generated from the supermatrix of the transcriptome-only dataset, the tree from the combined dataset recovered the free-living caddisfly families (Rhyacophilidae and Hydrobiosidae) as a monophyletic clade with tortoise-case makers (Glossosomatidae) as their sister group. Together they are reciprocally monophyletic with the tube-case makers (Phryganides). Four cluster likelihood mapping on the combined dataset recovered strong support for this arrangement with no evidence of bias in the permutation tests (Fig. S2).

**Figure 1.**
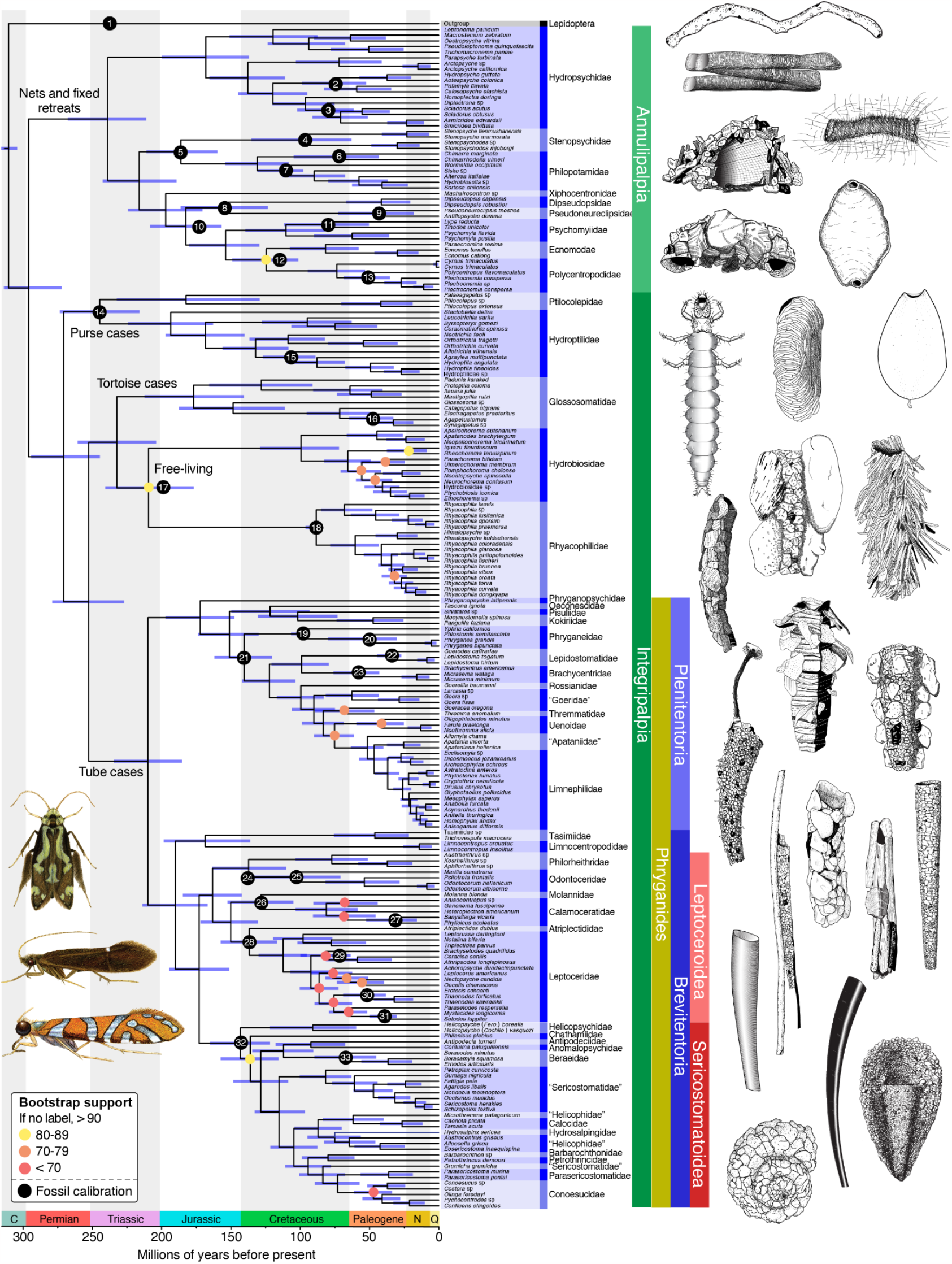
Dated phylogeny of Trichoptera from the combined targeted enrichment and transcriptome data set. Nodes with bootstrap support of less than 90 are labeled with yellow (80-89), orange (70-79), or red (<70) dots. The placement of fossil calibrations is indicated with numbered black dots. Selected larval caddisfly retreats and cases are shown on the right, and selected adult caddisflies are shown on the bottom left. Illustrations by Ralph Holzenthal, Julie Martinez, and Kristin Kuda.

### Molecular dating

We found that Trichoptera originated ~290 million years ago in the early Permian (Fig. 1). All extant higher groups within the order, however, originated in the Triassic, including Annulipalpia (fixed retreat makers), Hydroptiloidea (purse-case makers), Rhyacophiloidea (free-living and tortoise-case makers), and Phryganides (tube-case makers). Notably, tube-case making caddisflies within the infraorder Plenitentoria, the group that includes multiple lineages that construct their cases from flowering plant matter, originated in the mid-Jurassic ~170 million years ago.

### Ancestral state reconstruction and diversification analysis

We conducted ancestral state reconstructions on both the combined and transcriptome-only datasets to estimate the ancestral states for case-making and cocoon-making behavior in immature caddisflies. We found that the ancestral trichopteran was free-living and constructed a pupal dome (Fig. 2). We additionally recover three independent origins of portable case-making behavior on each of the branches leading to Hydroptilidae (purse-case makers), Glossosomatidae (tortoise-case makers), and Phryganides (tube-case makers).

**Figure 2.**
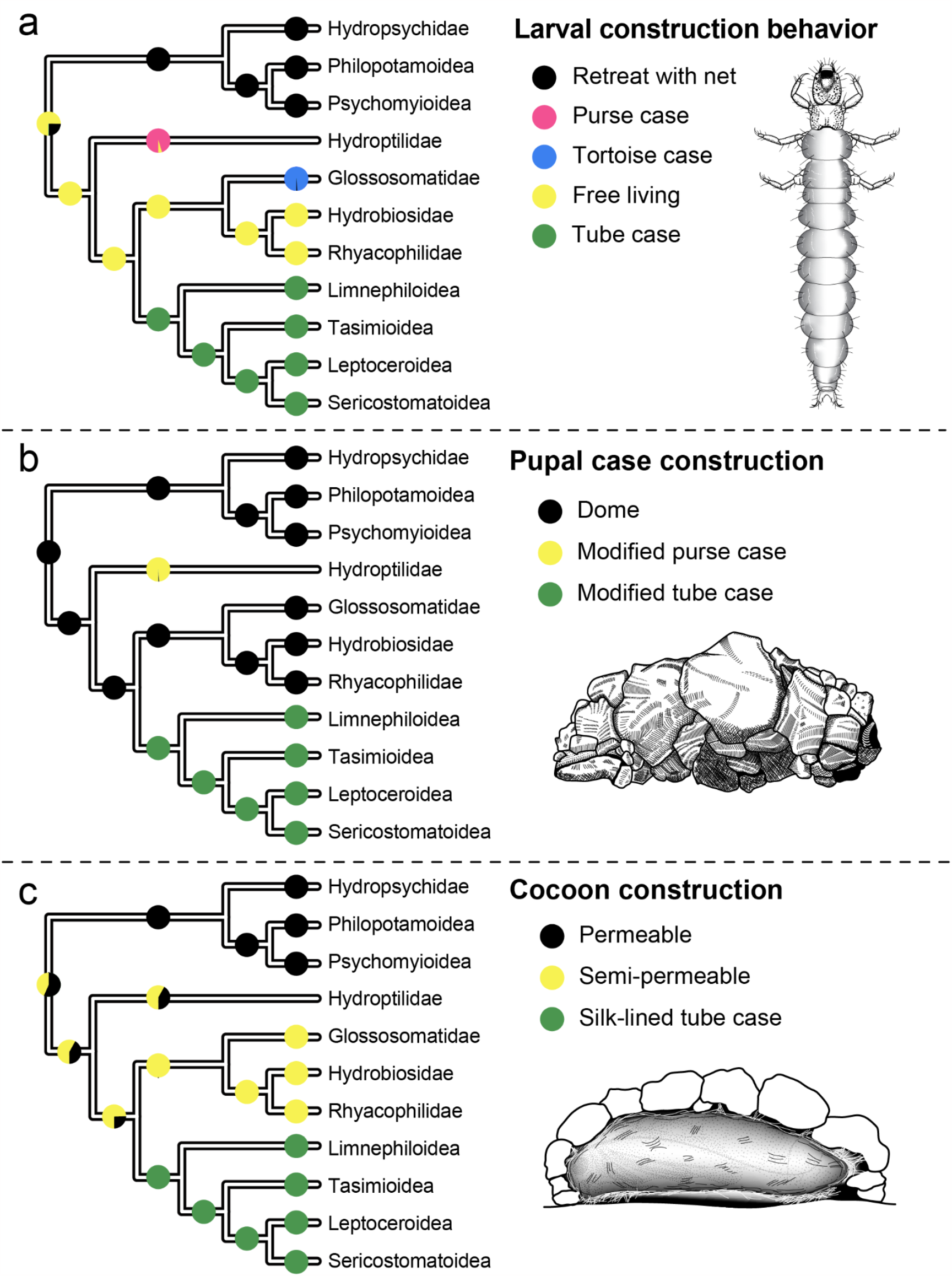
Results of ancestral state reconstruction (ASR) analyses, showing the ancestral states for (a) larval construction behavior, (b) pupal case construction, and (c) cocoon construction. Trees are simplified to major splits for visualization. Only three of the five character states in the pupal case construction ASR are present (with probability > 0.001) in the nodes visible in the simplified tree presented here. Illustrations by Ralph Holzenthal.

We recovered five shifts in diversification rate across the caddisfly tree using BAMM (Rabosky 2014) on the dated phylogeny from the combined dataset (Fig. 3). Three occurred during the Triassic and correlated with major groups: the suborder Annulipalpia or fixed-retreat makers; Hydroptilidae, the diverse family of microcaddisflies; and the lineage leading to the predatory free-living caddisfly families Hydrobiosidae and Rhyacophilidae. The other two shifts we recovered were nested within tube-case makers on the lineages leading to the diverse families Leptoceridae and Limnephilidae. We also generated a 16,688 taxon tree with stochastic polytomy resolution using Taxonomic Addition for Complete Trees (TACT) (Chang et al. 2020). This method allows for the addition of taxa for which there is no molecular information to a phylogeny based on the taxonomic classification of each species alone using birth-death sampling. On this tree, we recovered nearly 30 rate shifts, perhaps due to the large number of additional taxa (Fig. S3). While more shifts were recovered on this topology, many shifts coincided with those from the diversification analysis on the phylogeny from the combined dataset.

**Figure 3.**
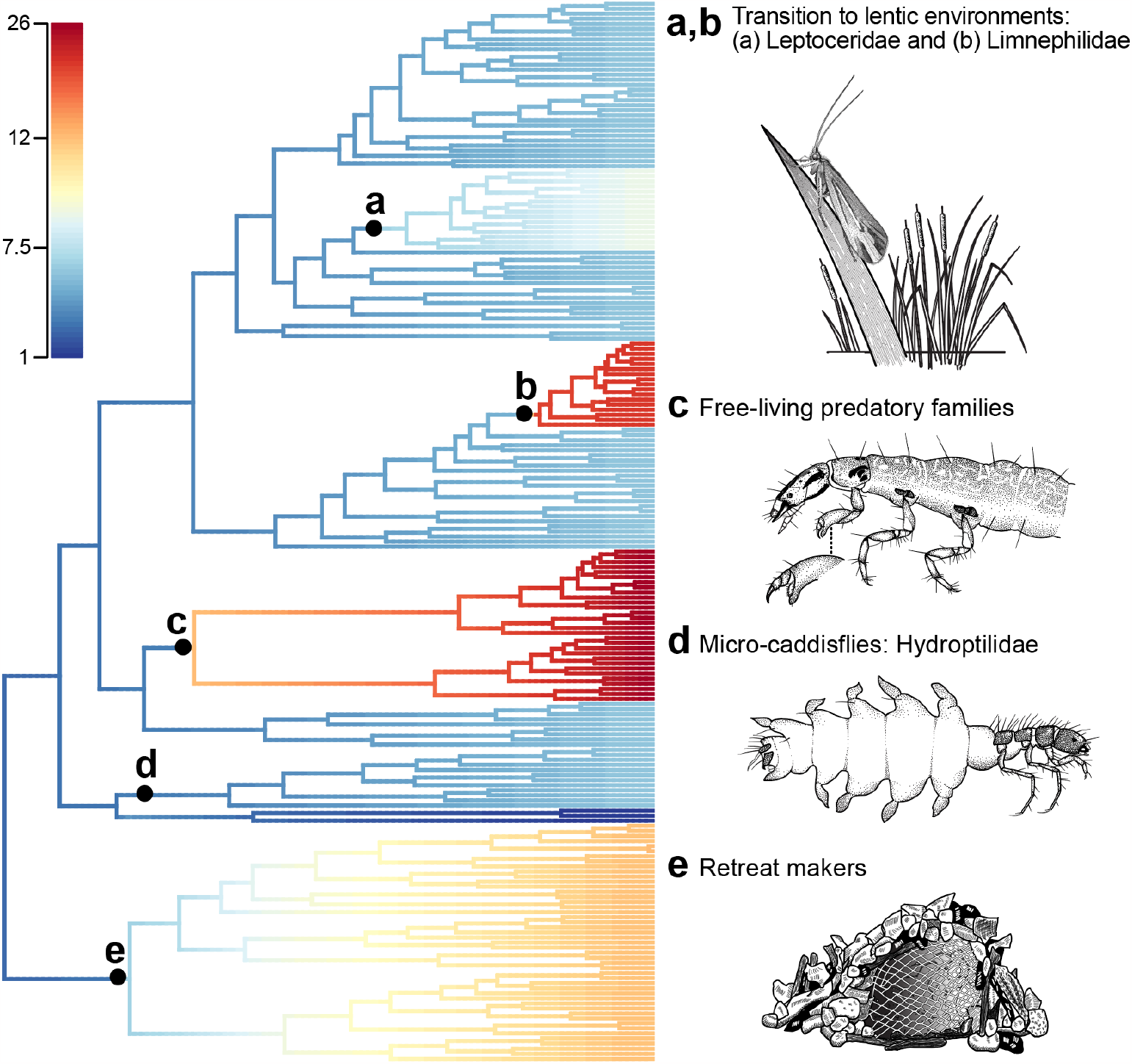
Diversification rates estimated on the Trichoptera phylogeny, corrected for unequal taxon sampling. Recovered rate shifts are shown in labeled black dots. Figure shows (a,b) two rate shifts that occurred in two tube-case making lineages that invaded lentic environments, (c) a shift in the lineage leading to predatory, free-living caddisfly families Rhyacophilidae and Hydrobiosidae, (d) a shift in the lineage leading to the major family of microcaddisflies, Hydroptilidae, and (e) a shift in the lineage leading to the suborder Annulipalpia. Illustrations by Ralph Holzenthal.

## Discussion

Trichoptera is the most diverse of the primarily aquatic insect orders and represents the second most diverse freshwater radiation of animals on the planet, surpassed perhaps only by the aquatic clade of true flies, Culicomorpha/Psychodomorpha (Malm et al. 2013). Their larvae use underwater adapted silk to extend their phenotypes in myriad forms, resulting in underwater architecture that provides camouflage and protection, aids in respiration, facilitates feeding, and stabilizes the stream beds they inhabit. These extended phenotypes are central to the diversification of the order. The first step toward a better understanding of their diversity is to contextualize the mode of their evolution with a robust phylogenetic tree. Here, we recovered a strongly supported phylogeny for 206 species, representing 48 of the 52 extant families (Fig. 1). Further, we find that portable case-making, responsible for more than half of the diversity of described caddisfly species (Morse 2009), evolved on three separate occasions from an ancestral free-living form (Fig. 2). However, despite repeated evolution of portable cases, diversification shifts occurred later, corresponding to other ecological innovations, some only made possible by the earlier use of a portable case (Fig. 3).

The relationships among suborders within Trichoptera have been a point of conflict since the first evolutionary trees were drawn (Martynov 1924; Milne and Milne 1939; Ross 1956), pre-dating widespread use of Hennigian systematics. Since then, the primary sources of incongruence among phylogenetic estimates were the relationships among the monophyletic tube-case makers (Phryganides), the monophyletic fixed-retreat makers (Annulipalpia), and the five remaining families, the free-living families (Rhyacophilidae and Hydrobiosidae), the tortoise-case makers (Glossosomatidae), and the two families of micro-caddisflies (Ptilocolepidae and Hydroptilidae). Multiple different arrangements of the relationships among these families were posited, using both morphology and molecular data (Ross 1967; Weaver III 1984; Wiggins and Wichard 1989; Frania and Wiggins 1997; Kjer et al. 2001, 2016; Ivanov 2002; Malm et al. 2013; Thomas et al. 2020; Ge et al. 2023). Thomas et al. (2020) analyzed five nuclear genes and found strong support for the cocoon-making families as a grade leading to the tube-case makers, resulting in their reclassification into the suborder Integripalpia. However, the relationships among the families of cocoon-makers were not well-supported, preventing the interpretation of ancestral states. More recently, an analysis of mitochondrial genomes recovered a similar arrangement with the exception that the microcaddisflies (Hydroptilidae) were sister to the rest of caddisflies (Ge et al. 2023). Here, we estimated a phylogeny with strong support for the grouping of the families of cocoon-makers with the tube-case making families (Phryganides). Further, we find strong support for a clade that includes tortoise-case makers (Glossosomatidae) and the two free-living families (Rhyacophilidae and Hydrobiosidae). Though this clade had not previously been recovered with molecular data, morphological experts have long contested that the pupal forms of all three families suggested a common origin (Wiggins 2005), and that the tortoise cases of Glossosomatidae likely evolved as “precocious pupation behavior.” Our results are consistent with this interpretation. However, while the monophyly of this clade was strongly supported, the relationships among families within the clade differed among data sets and tree-estimation strategies. Four-cluster likelihood mapping revealed a bias in the maximum likelihood analysis of the concatenated supermatrix of transcriptomes, indicating that the relationships recovered in the ASTRAL-III analysis and in the maximum likelihood analysis of the combined dataset, which place Hydrobiosidae as sister to Rhyacophilidae are the most reliable (Fig. S1).

Caddisflies began to diversify in the Permian, about 295 million years ago. Our results suggest that the ancestor of all extant caddisflies roamed the benthos of the oxygen-saturated waters of rivers and streams as a free-living larva, without a portable case or fixed retreat. Like its amphiesmenopteran ancestors, the larva produced silk from labial glands. As it approached pupation, the final instar larva constructed a protective pupal structure of small mineral fragments or other substrates and enclosed itself within the structure in a permeable or semi-permeable silken cocoon. As caddisflies further evolved and diversified, selection favored the precocious retention of the pupal structure by earlier instars and increasingly complex larval structures. In the suborder Annulipalpia, larvae affixed various materials from the stream bed into retreats, offering protection. Eventually, their silk use diversified across the suborder to include capture nets and sophisticated filtration systems associated with their retreats to capture suspended organic particles. In the suborder Integripalpia, due to the efficacy of the case in providing physical protection, camouflage, and more efficient respiration, three different lineages co-opted these precocious pupal structures into portable cases, eventually resulting in purse cases (only in the final instar larvae in Hydroptilidae and Ptilocolepidae), tortoise-cases (Glossosomatidae), and tube-cases (Phryganides). However, the primary species diversification events in case-makers did not coincide with the evolution of the case, but with other ecological innovations. For example, bursts in diversification occurred in the lineages leading to the families Leptoceridae and Limnephilidae coinciding with their incorporation of plant matter for case-making. Simultaneously, this allowed them to invade slow-moving and standing waters facilitated by the respiratory efficiency of tube cases to enhance the flow dynamics of oxygenated water across the larval abdomen and gills (Wiggins 1996). Since that time, caddisflies have continued to diversify in their respective habitats, eventually becoming one of the dominant animals in freshwater ecosystems.

## Materials and Methods

### Taxon sampling

We generated transcriptome data for 50 new species of caddisflies and targeted enrichment data for an additional 158 individuals representing 155 species. We also incorporated one published trichopteran genome into our analyses, resulting in a dataset that comprises 209 ingroup taxa, representing 206 trichopteran species, 174 genera, and 48 of 52 extant families. We used 18 outgroup taxa, including 17 lepidopterans and one coleopteran harvested from publicly available transcriptomes and genomes, for a total count of 227 individuals.

### Transcriptome sequencing

We generated 50 new caddisfly transcriptomes representing 34 families and all of the major clades within the order. Specimens were collected alive and preserved in RNAlater. Specimens were kept cool until RNA extraction. Libraries were prepared and sequenced at BGI-Shenzhen on an Illumina HiSeq 2000. Full details are given in Supplemental Note 1.

### Anchored hybrid enrichment probe design and sequencing

Despite extensive efforts to sample and sequence a diverse sample of caddisflies using transcriptomes, our sampling was limited by the species we could collect and identify while alive, before destructively preserving them in RNAlater. To expand our taxon sampling to include museum specimens, we used anchored hybrid enrichment (AHE; (Lemmon et al. 2012). First, we used assembled transcriptomes from 15 species spread throughout the order to identify exonic regions of high conservation, flanked by regions that were more variable for phylogenetic inference. Using this method, we identified ~900 exons suitable for phylogenetic inference (Supplemental Note 2).

DNA was extracted from museum specimens using the G-biosciences Q-amp extraction kit. Extracted DNA was sonicated using the Covaris Ultrasonicator to a size distribution of 150-400bp. Libraries were prepared following Lemmon et al. (Lemmon et al. 2012) and indexed with single 8-bp indexes chosen to be different at least two sites. Libraries were then combined in pools of ~16 samples and enriched using the aforementioned enrichment kit. After qPCR-based QC, enriched libraries were sequenced on two lanes of an Illumina HiSeq 2500 sequencer with a paired-end 150bp sequencing protocol and onboard cluster generation.

### Transcriptome-only analyses

We followed the 1KITE transcriptome phylogenetic pipeline as described in Wipfler et al. (2019). Briefly, transcriptomes were assembled in SOAPdenovo-Trans (Xie et al. 2014). Following transcriptome assembly, we used a set of core orthologs using the genomes from *Tribolium castaneum, Danaus plexippus*, and *Bombyx mori* (Kawahara et al. 2019) to predict orthologs using Orthograph (Petersen et al. 2017). We compiled single-copy orthologs for each trichopteran species and lepidopteran outgroup into individual ortholog files. We aligned the amino acid alignments with MAFFT v.7.475 (Katoh and Standley 2013) using the L-INS-i algorithm. We then evaluated the alignments for outliers using the pipeline described in Misof et al. (2014). Following alignment, we used Aliscore v.02.2 and ALICUT v.2.3 to identify and remove portions of the alignment that were indistinguishable from random data (Misof and Misof 2009). Because some gene alignments may not be phylogenetically informative, we used MARE v.1.2-rc (Meyer et al. 2011) to remove genes with 0 information content. Finally, we generated a concatenated alignment supermatrix using FASconCAT v.1.0 (Kück and Meusemann 2010).

We generated phylogenetic trees using two methods: (1) a maximum likelihood (ML) tree generated from the concatenated supermatrix using partitioned models of molecular evolution and (2) a multispecies coalescent (MSC) tree using ASTRAL-III (Zhang et al. 2018). ASTRAL-III requires individual gene trees as input, thus we generated individual gene trees using maximum likelihood with IQ-TREE v. 1.6 (Nguyen et al. 2015). For each individual gene tree, we selected models of molecular evolution using ModelFinder (-MFP option) as implemented in IQ-TREE v. 1.6 and performed 25 separate ML tree searches for each gene. We selected the best tree for each gene and concatenated them for input into ASTRAL-III. We then ran ASTRAL-III using default settings. For the concatenated supermatrix analysis, we first selected an optimal partitioning scheme with PartitionFinder 2 (Lanfear et al. 2017) followed by the selection of protein models for each metapartition using ModelFinder as implemented in IQtree v. 1.6 (-MFP option). Following the selection of models of molecular evolution, we conducted 20 individual ML tree searches on the partitioned supermatrix (10 with a random starting tree and 10 with an IQtree-generated parsimony starting tree).

To further evaluate relationships that were incongruent between the ML analysis on the concatenated supermatrix and the ASTRAL-III analysis, we performed Four-cluster Likelihood Mapping (FcLM) on the concatenated supermatrix using IQ-TREE v. 1.6. In addition to the FcLM, we performed three series of permutation tests, a strategy previously used by Misof et al. (2014), to evaluate potential confounding signal arising from among lineage heterogeneity and the distribution of missing data (Supplementary Note 3).

### Targeted enrichment analyses

Raw read pairs passing the CASAVA high-chastity filter were merged following Rokyta et al. (2012) and library adapters were trimmed. We then followed the pipeline established by Breinholt et al. (2018) for assembly and ortholog prediction. In short, we assembled each locus using an iterative-baited assembly process, where reads are mapped to reference loci and then assembled in local assemblies with Bridger (Chang et al. 2015). We determined orthology by searching each locus against the reference genome using BLAST (Altschul et al. 1990). If the locus generated hits to the reference genome in more than one contig or if the reciprocal BLAST hit matched a different locus from the species sampled, the gene was removed.

We then translated the alignments by predicting open reading frames in each locus and choosing the reading frame that included no stop codons in any taxon.

### Combined analyses

While the targeted enrichment probes were designed from the trichopteran transcriptomes, only portions of the original transcriptome assemblies were included in the final ortholog set. Thus, when combining datasets, we first needed to identify loci that overlapped in both datasets. To do this, we used reciprocal blastp searches to identify putative overlapping orthologs. We aligned each ortholog with MAFFT v. 7.475 using the linsi algorithm (Katoh and Standley 2013). Following ortholog prediction, we generated trees for each gene. In some cases, taxa derived from transcriptome data and taxa derived from targeted enrichment data formed separate clades, indicating a clear orthology mismatch. Each tree and corresponding alignment were inspected by hand, and such ortholog misassignments were removed from further analysis.

We concatenated the individual genes of the combined data into a supermatrix using FASconCAT v.1.0 (Kück and Meusemann 2010). We then selected a partitioning scheme and models in IQ-TREE v.1.6 using the -MERGE+MFP option (Nguyen et al. 2015; Kalyaanamoorthy et al. 2017). Using this model, we conducted 30 individual tree searches in IQ-TREE (15 with a random starting tree and 15 with a parsimony starting tree) and selected the best tree for further analysis. We used the -bb option in IQ-TREE with 1000 replicates to estimate ultrafast bootstrap support and used the -bnni option to correct for potential overestimation of support in the presence of model violations. We also repeated the FcLM analyses on the combined dataset to evaluate the source of bias or signal regarding the relationships among the families of free-living (Rhyacophilidae and Hydrobiosidae) and tortoise case-making (Glossosomatidae) caddisflies (Supplementary note 3).

### Molecular dating, ancestral state reconstructions, and diversification analyses

We selected 33 fossils (32 Trichoptera and one Lepidoptera) to calibrate our tree. We then performed two dating analyses in MCMCtree (Yang 2007), one with uniform priors and another with truncated cauchy priors. The general methods follow Kawahara et al. (2019). In short, we estimated Hessian matrices on the aligned amino acids using the LG substitution model and five rate categories. We used CODEML to estimate empirical base frequencies. We then used the estimated Hessian matrix with a fixed topology from the best maximum likelihood tree (combined transcriptome and targeted enrichment) as input to MCMCtree. We ran four analyses for each set of node calibration priors (uniform and cauchy). We additionally ran the time priors for each analysis without the dataset to ascertain the informativeness of the priors (Fig. S4-5). We checked each run for convergence using Tracer v. 1.7 (Rambaut et al. 2018) and by plotting the posterior mean ages and the lower and upper bounds for the 95% credibility intervals for each of the runs against each other (Fig S5-6). Converged runs were then combined.

Ancestral state reconstructions (ASRs) for larval and pupal morphology were performed on the ML trees generated from the transcriptome-only and the combined datasets. Stochastic character mapping was used to conduct all ASRs, with the ‘make.simmap’ command in the R package Phytools v07-70 (Revell 2012). One thousand stochastic maps were generated for each ASR using a symmetric-rate model, which assumes pairs of forward and reverse character state transitions have equal rates. Character matrices were generated for the following three discrete characters: 1. Larval construction behavior, coded with five states (Fixed retreat with net, purse case, tortoise case, no structure [i.e., free-living], tube case); 2. Pupal case construction, coded with five states (Dome-shaped enclosure, fixed silken larval tube case, modified purse case, newly constructed rigid tube case, modified larval tube case); 3. Cocoon construction, coded with three states (Semi-permeable cocoon, permeable cocoon, silk-lined tube case).

We used two strategies to estimate diversification rates in BAMM (Rabosky 2014). In the first, we estimated diversification rates using the ultrametric tree derived from the MCMCtree dating analysis (Fig. 1) and accounted for unequal taxon sampling using lineage-specific sample proportions at the family level. In the second, we used the stochastic polytomy resolution method implemented in “Taxonomic Addition for Complete Trees” (TACT) (Chang et al. 2020) to add unsampled taxa to the tree. We used the complete known taxonomy for Trichoptera as recorded in the Trichoptera World Checklist (Morse 2009), resulting in a tree with 16,688 tips. For both datasets, we ran four separate diversification analyses in BAMM for 10 million generations. For each dataset, we used BAMMtools in R (Rabosky et al. 2014) to plot the log-likelihood against the generation number to assess burn-in and subsequently removed the first 20% of each run where the log-likelihood reached an asymptote. We then estimated the ESS using the R package coda. For each analysis, we then generated both the credible shift set and the maximum shift credibility in BAMMtools and compared the output from the four runs on each dataset to assess convergence.

## Contributions

P.B.F., B.M., and X.Z conceived the study. P.B.F, M.E., J.B., J.A.T., S.S., D.P., Y.L., O.N., C.M., L.P., A.D., B.M., E.M.L., A.L., J.C.M. and X.Z. contributed analyses. P.B.F., B.M., S.P., and X.Z. acquired funding. P.B.F., M.E., J.B., J.A.T., S.S., A.Y.K., D.P., O.N., C.M., L.P., A.D., B.M., E.M.L, A.L., J.C.M. and X.Z. performed data assembly. P.B.F. and X.Z. were responsible for project administration. P.B.F., R.W.H., A.Y.K., C.R.N., S.P., and X.Z. acquired samples. P.B.F., R.W.H., M.E., S.S., D.P., and S.H. performed visualizations. P.B.F., R.W.H., M.E., and S.H. wrote the original draft of the article. All authors reviewed and edited subsequent drafts.

## Acknowledgments

Analyses were performed on the Smithsonian High Performance Cluster (SI/HPC), the BYU Office of Research Computing Supercomputer, the LIB High Performance Cluster (LIB HPC), and the HiPerGator computing cluster at the University of Florida. We thank Julie Martinez for the illustrations of adult caddisflies showcased in Figure 1. We thank the 1KITE consortium from which this paper emerged, especially Karen Meusemann and Karl Kjer for their support and advice. Sequencing and assembly of the 1KITE transcriptomes were funded by BGI through support to China National GeneBank.

