## supplemental_information for "Phylogenomics recovers multiple origins of portable case-making in caddisflies (Insecta: Trichoptera), the world’s most common underwater architects"

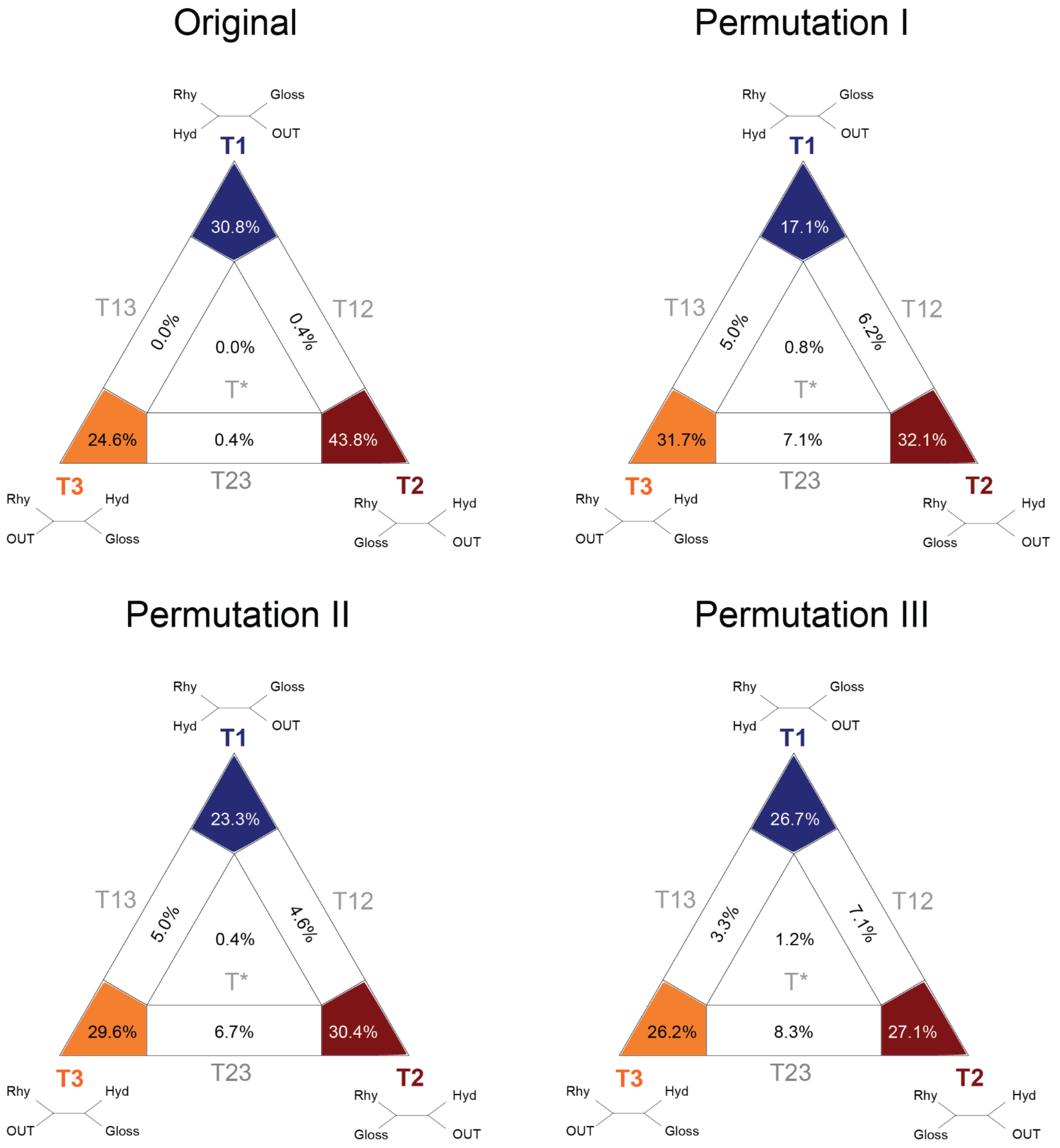


**Fig. S1.** Four cluster likelihood mapping (FcLM) of the relationships between free-living (Rhyacophilidae (Rhy) and Hydrobiosidae (Hyd)) and tortoise-case making (Glossosomatidae (Gloss)) families for the transcriptome only dataset. Permutation tests were conducted to evaluate the effect of bias on the dataset. Though the FcLM on the original dataset shows more support for hypothesis, T2, Permutation I shows that the transcriptome only dataset is biased toward T2 and T3 due to among-lineage heterogeneity and non-random distribution of missing data.

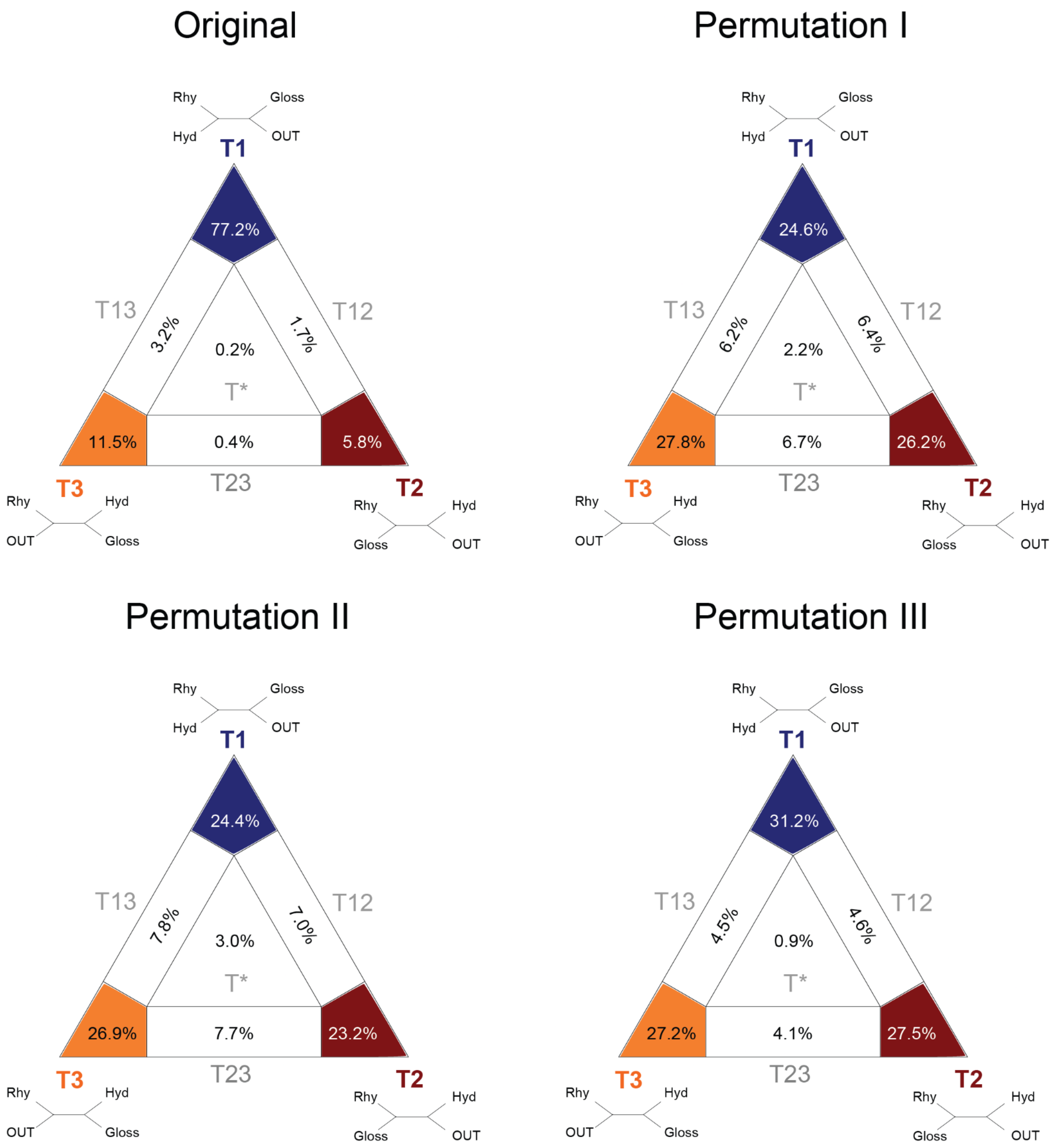


**Fig. S2.** Four cluster likelihood mapping (FcLM) of the relationships between free-living (Rhyacophilidae (Rhy) and Hydrobiosidae (Hyd)) and tortoise-case making (Glossosomatidae (Gloss)) families for the combined dataset. Permutation tests were conducted to evaluate the effect of bias on the dataset. In this dataset, T1 is the most strongly supported hypothesis with no effect of bias found in the Permutation tests.

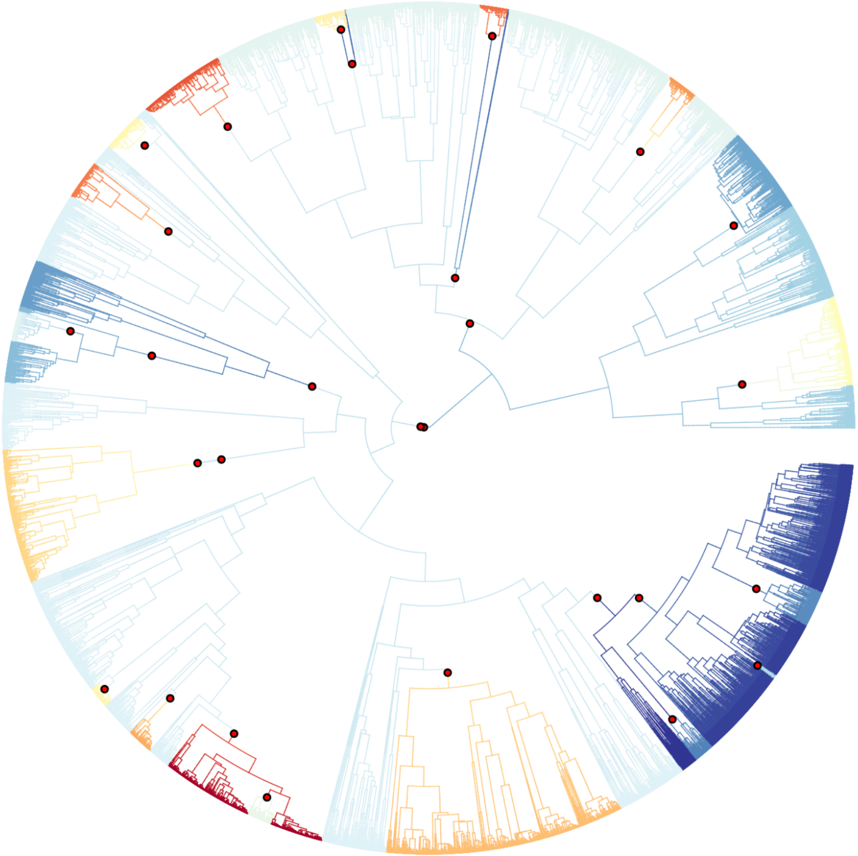

**Fig. S3.** Diversification results from tree supplemented with stochastic polytomy resolution using Taxonomic Addition for Complete Tree (TACT) to place unsampled taxa. Red dots indicate diversification rate shifts.

**Supplemental Note 1**

**RNA extraction and sequencing**

The newly sequenced specimens for this study were collected as either larvae or adults, preserved in RNAlater (Qiagen, Maryland, USA), and stored at +4 °C or -80 °C until further processing. Whenever possible, specimen vouchers for the newly-generated transcriptomes were kept at BYU. For small specimens, the entire body was ground with a pestle and used for extraction of total RNA. For large-bodied specimens, only a piece of the thoracic tissue (usually flight muscles) was used. Total RNA extractions, mRNA isolation, fragmentation, and cDNA library construction were performed using the protocols of Misof et al. (2014) and Peters et al. (2017). Paired-end sequencing of the transcriptome libraries was conducted through the Beijing Genome Institute (BGI, Shenzhen, Guangdong, China) on HiSeq 2000 or 2500 platforms (Illumina, San Diego, CA, USA) with read lengths of either 90 bp (for libraries prepared with the TruSeq kit) or 150 bp (for libraries not prepared with the TruSeq kit). Approximately 2.5 gigabases of raw data per library were retained.

**Supplemental Note 2**

**Anchored hybrid enrichment probe design, DNA extraction, and sequencing**

We used assembled transcriptomes from 15 species spread throughout the order to identify exonic regions of high conservation, flanked by regions that were more variable for phylogenetic inference. Using this method, we identified ~900 exons suitable for phylogenetic inference. The probes were designed in collaboration with the Center for Anchored Phylogenomics. Target AHE loci in common among Trichoptera and other Holometabola were identified by scanning fifteen trichopteran 1Kite transcriptomes for the Lepidoptera AHE loci identified by Breinholt et al. (2018). The transcripts identified were aligned in MAFFT v7.023b (2013) by target locus, then trimmed to well-aligned regions, and finally manually inspected in Geneious R9, (Biomatters Ltd., Kearse et al. 2012). A total of 960 target loci (averaging 232bp in length) remained after masking/removing regions identified to be repetitive using kmer distribution profiling (see Hamilton et al. 2016 for details). For each of these loci, probes were tiled uniformly across each of the sixteen reference sequences at 4.2x coverage to produce 57094 probes. Probes were produced by Agilent as a SureSelect XP kit. DNA was extracted from museum specimens using the G-biosciences Q-amp extraction kit. Extracted DNA was sonicated using the Covaris Ultrasonicator to a size distribution of 150-400bp. Libraries were then prepared at the Florida State Center for Anchored Phylogenomics following Lemmon et al. (2012) and indexed with single 8-bp indexes chosen to be different at least at two sites. Libraries were then combined in pools of ~16 samples and enriched using the aforementioned enrichment kit. After qPCR-based QC, enriched libraries were sequenced on two lanes of an Illumina HiSeq 2500 sequencer with a paired-end 150bp sequencing protocol and onboard cluster generation (the raw data totaled 88 Gb).

**Supplemental Note 3**

**Topology tests**

We supplemented our bootstrap support values with another statistical approach, Four-cluster Likelihood Mapping (FcLM) (Strimmer & Haeseler, 1997), to evaluate support for alternative relationships. FcLM only addresses single splits in a tree. Therefore, this approach enables the identification of hidden signal for single relationships that may not be seen in ML trees. We tested the potential sister group relationship of Rhyacophilidae and Hydrobiosidae, which was recovered in the ML analyses of the combined dataset but not in the transcriptome-only dataset; that latter analysis instead recovered Rhyacophilidae as sister to Glossosomatidae.

To test this hypothesis about the phylogenetic placement of Rhyacophilidae, we defined four groups of interest: Rhyacophilidae (Rhy), Hydrobiosidae (Hyd), Glossosomatidae (Gloss), and outgroup taxa (OUT). We additionally checked for confounding signal due to among-lineage heterogeneity, non-random substitution processes and/or distribution of missing data by generating permuted datasets that lacked phylogenetic signal. In permutation I, all phylogenetic signal was removed. In permutation II, compositional heterogeneity was removed by randomly drawing amino acids using the frequencies of the LG substitution matrix, but the distribution of missing data was left untouched. Permutation III included the features of permutation II, but also had random distribution of missing data (i.e. no phylogenetic signal, homogeneous composition, and missing data randomly distributed). See Sann et al. (2018) and the Supplementary Information in Misof et al. (2014) for additional information on this methodology.

FcLM analyses were performed on both the transcriptome-only and the combined datasets using IQ-TREE version 1.6.beta4. The LG substitution model was used for each partition. Proportions of quartets that mapped into respective areas are provided in a 2D simplex graph (Fig. S1-2).

For both FcLM analyses, the three possible unambiguous topologies were:
T1: Rhy, Hyd | Gloss, OUT
T2: Rhy, Gloss | Hyd, OUT
T3: Rhy, OUT | Hyd, Gloss

The ML tree from the transcriptome-only dataset supports a sister group relationship of Rhyacophilidae and Glossosomatidae, and the results of the corresponding non-permuted FcLM analysis also favor this topology with a slight plurality of quartets (T2: 43.8%). However, permutation test I revealed that among-lineage heterogeneity and non-randomly distribution of missing data most likely biased the support for the sister group relationship of Rhyacophilidae and Glossosomatidae in the original (non-permuted) dataset and the ML tree reconstruction. Since the results of permutation test II are very similar to that of permutation test I, possible impact from confounding signal might rather be caused by non-random distribution of missing data than among-lineage heterogeneity. That means we cannot exclude the possibility that the ML tree's support for a monophyletic Rhyacophilidae+Glossosomatidae is biased from confounding signal, while other signal coming from potentially phylogenetic information might be overruled by confounding signal.

The ML tree from the combined dataset supports a sister group relationship of Rhyacophilidae and Hydrobiosidae, and the results of the corresponding non-permuted FcLM analysis strongly favor this topology with a majority of quartets (T1: 77.2%). The relatively even distribution of quartets in the permuted analyses (Fig. S2) indicates that this support for Rhyacophilidae + Hydrobiosidae cannot be explained by confounding signal. Therefore, we consider the placement of Rhyacophilidae as sister to Hydrobiosidae to be robust and not biased.

**Fig. S4 Density plots comparing time priors against posteriors for cauchy priors**

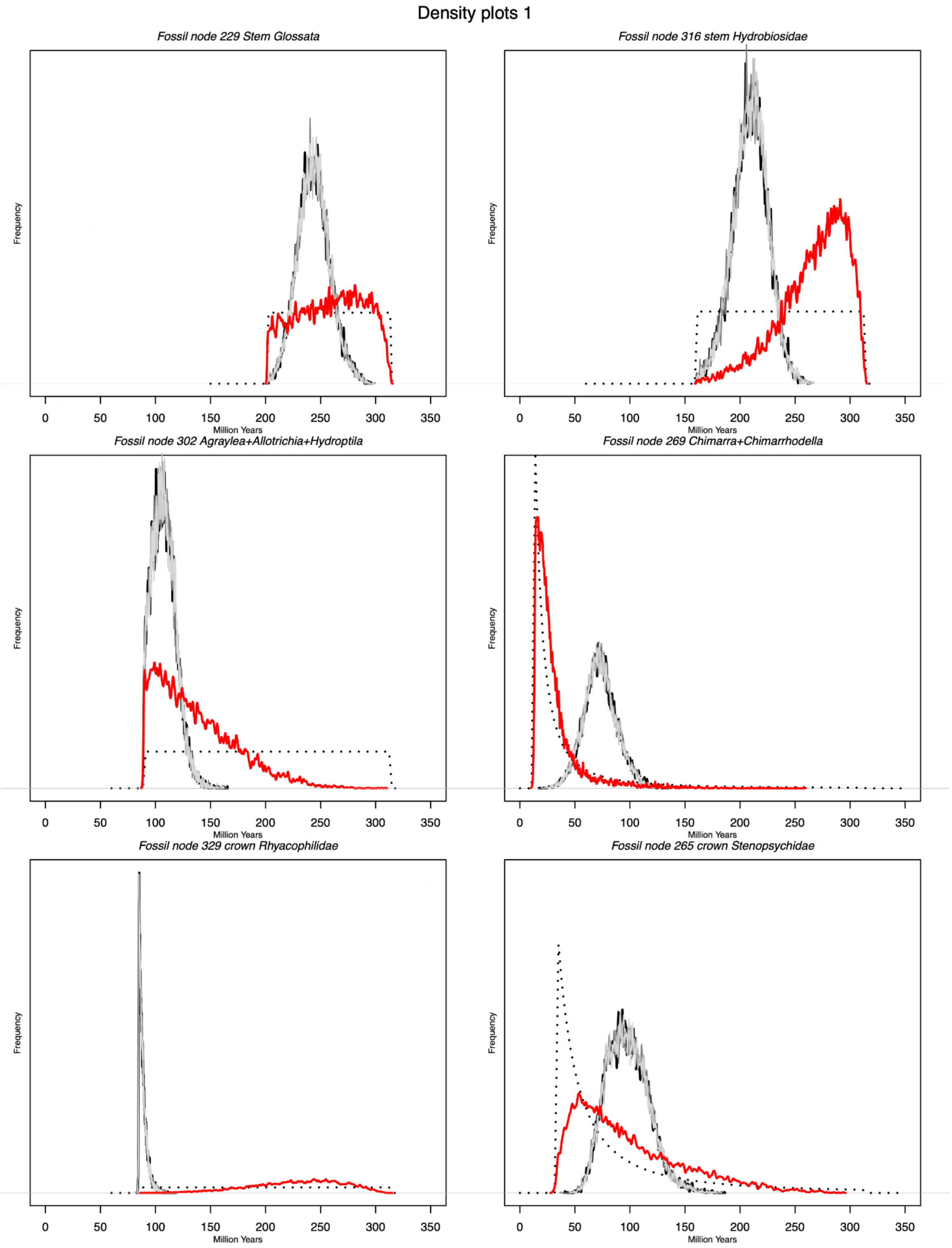

### Density plots 2

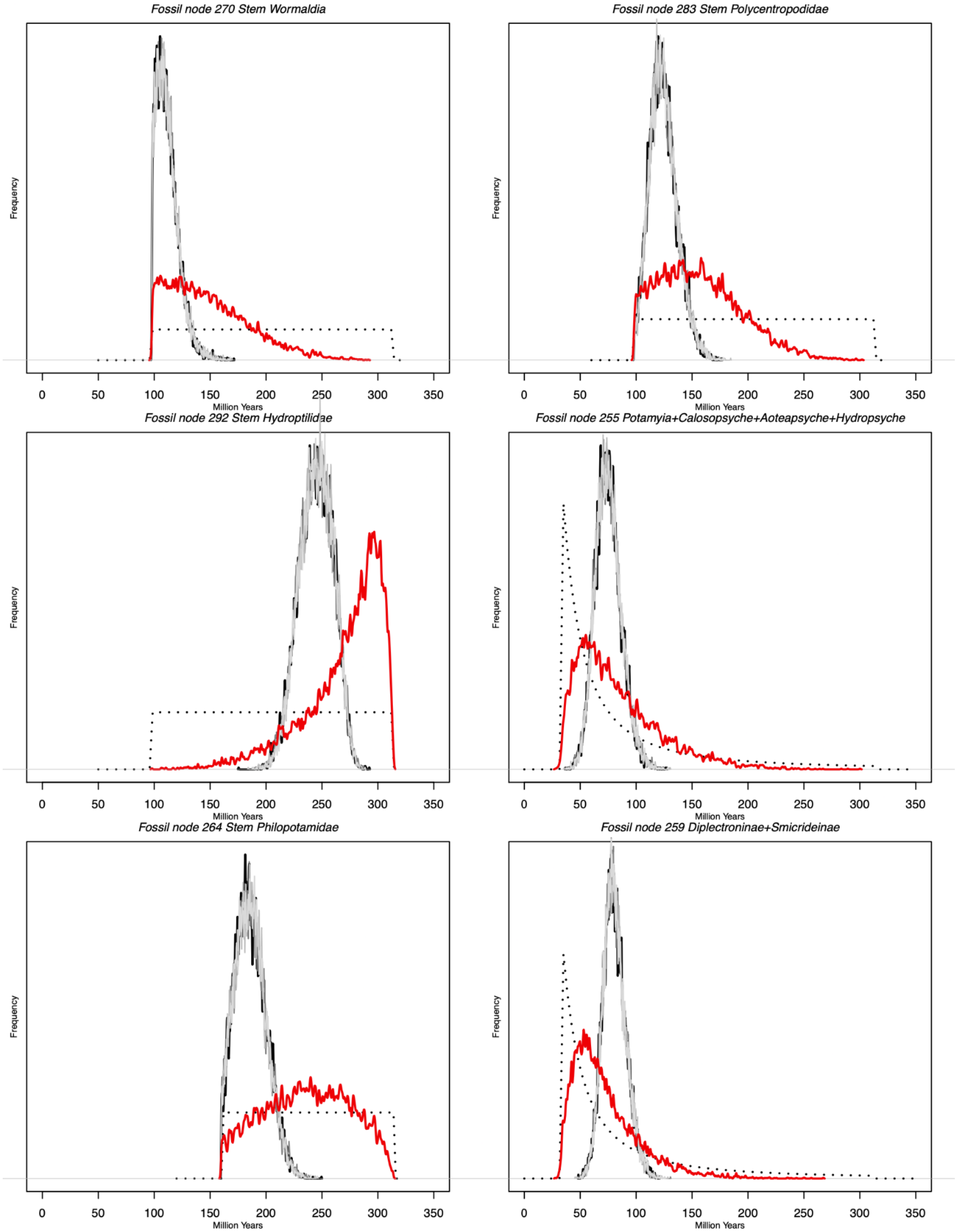

### Density plots 3

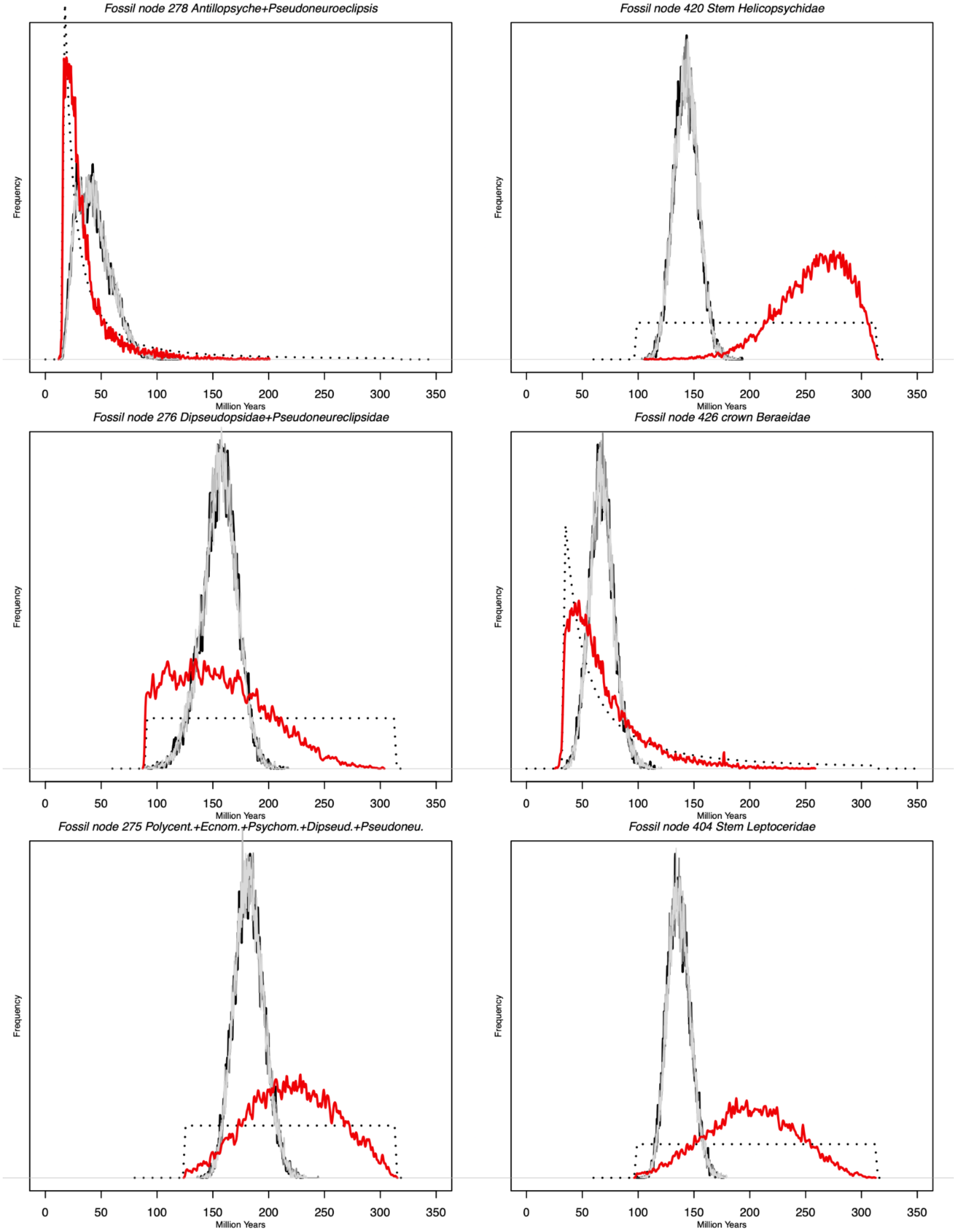

#### Density plots 4

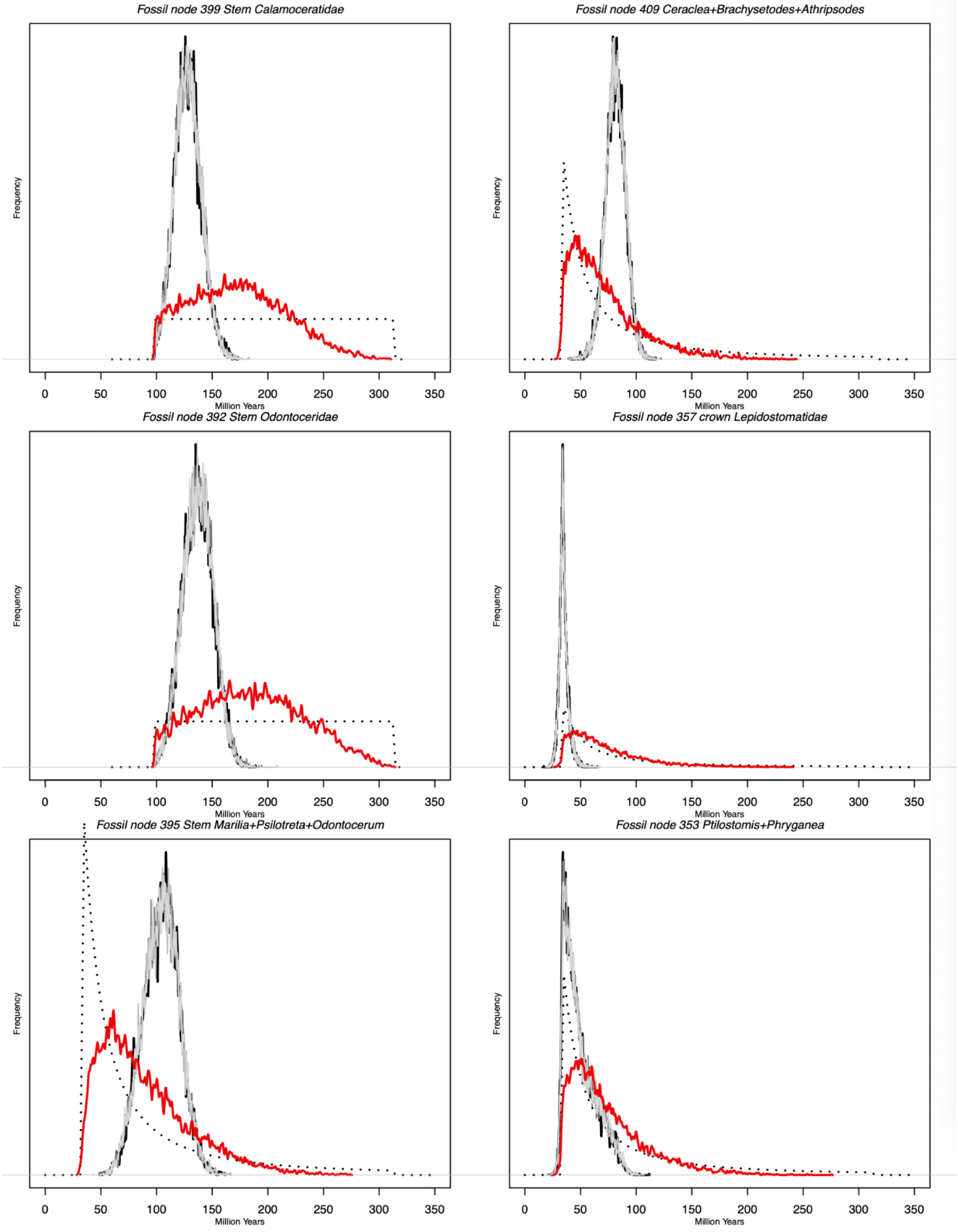

### Density plots 5

*Fossil node 359 crown Brachycentridae*

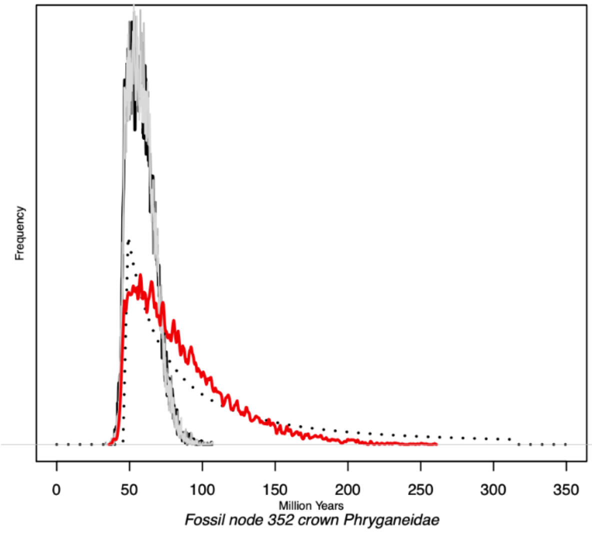


*Fossil node 419 Setodes+Mystacides*

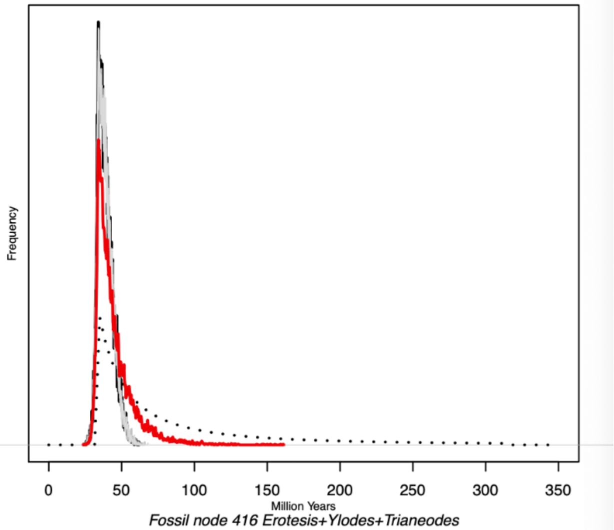


*Fossil node 352 crown Phryganeidae*

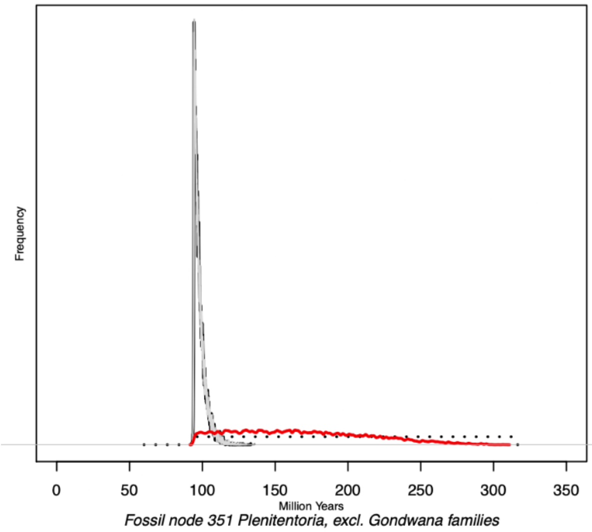


*Fossil node 416 Erotesis+Ylodes+Trianeodes*

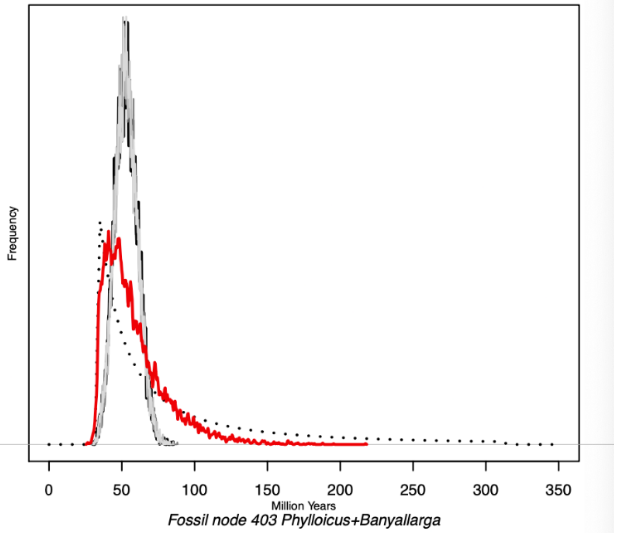


*Fossil node 351 Plenitentoria, excl. Gondwana families*

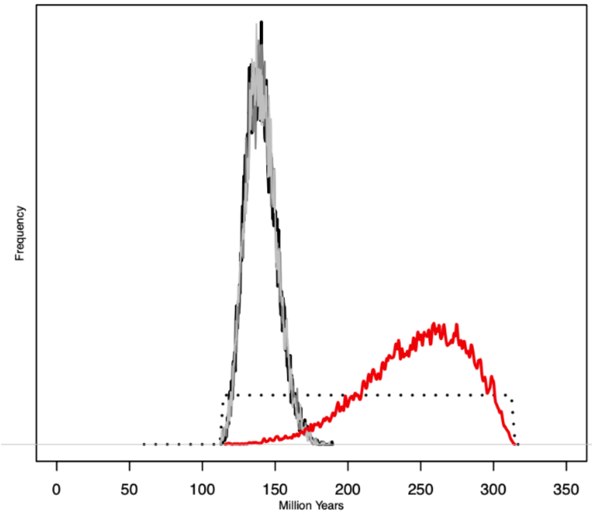


*Fossil node 403 Phylloicus+Banyallarga*

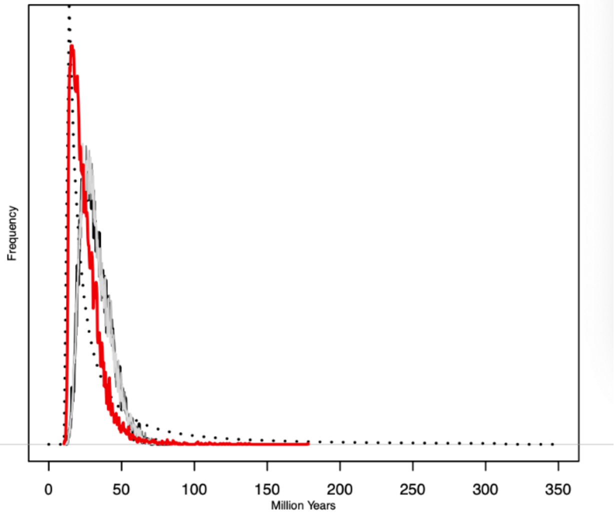

### Density plots 6

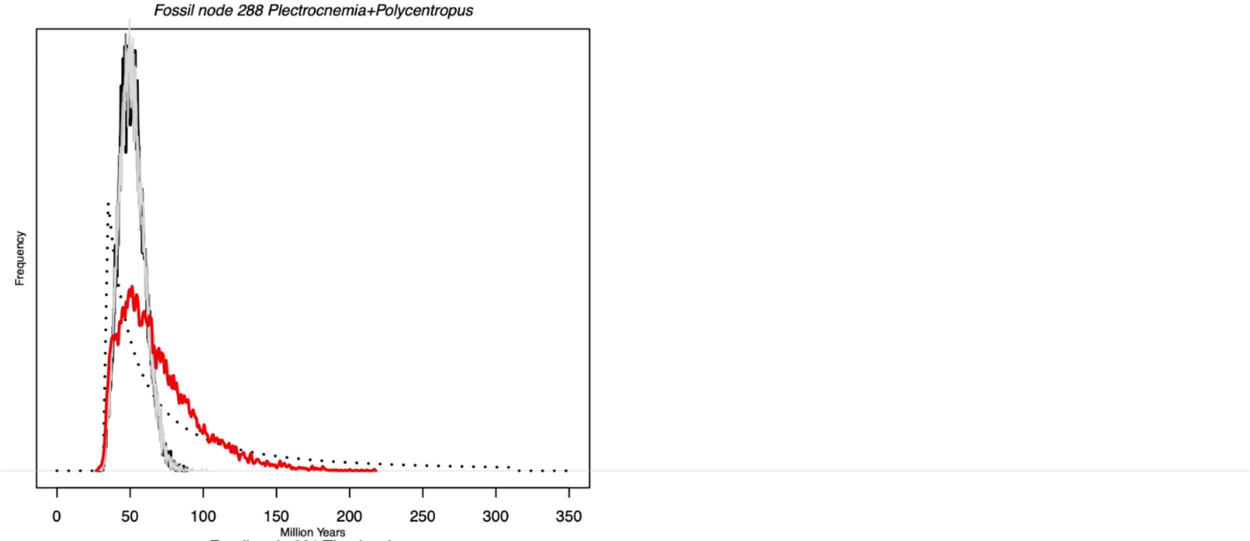


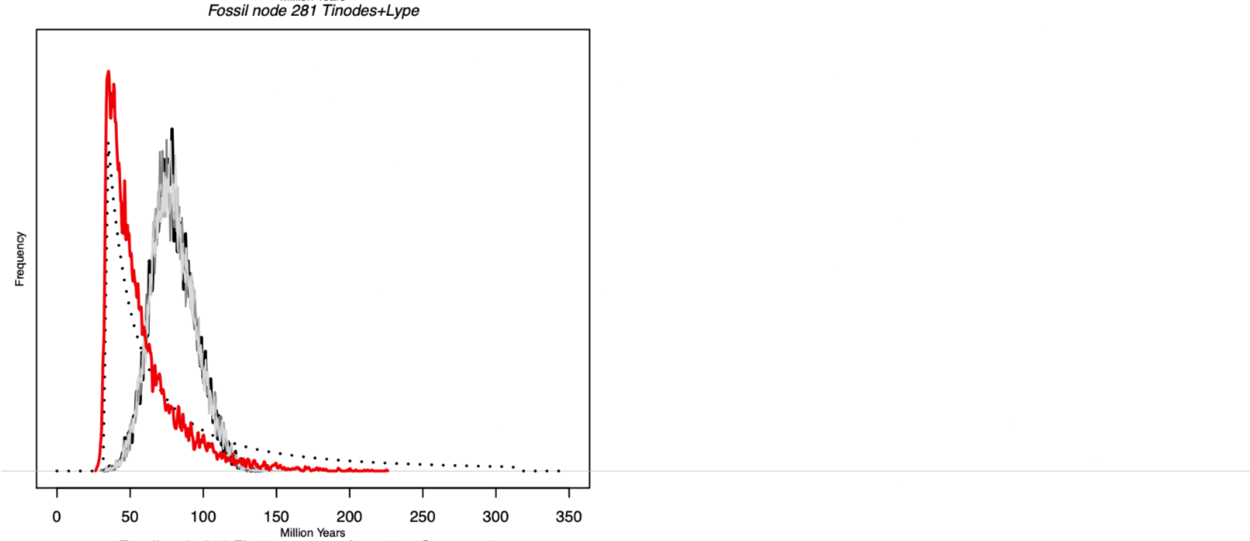


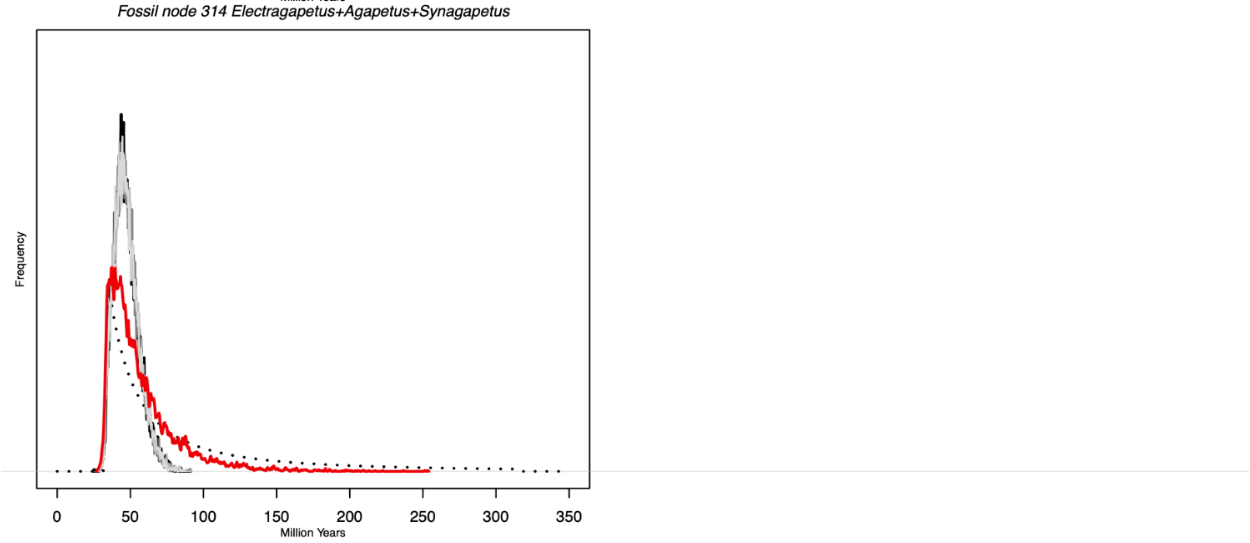

#### Fig. S5  Density plots comparing time priors against posteriors for uniform priors

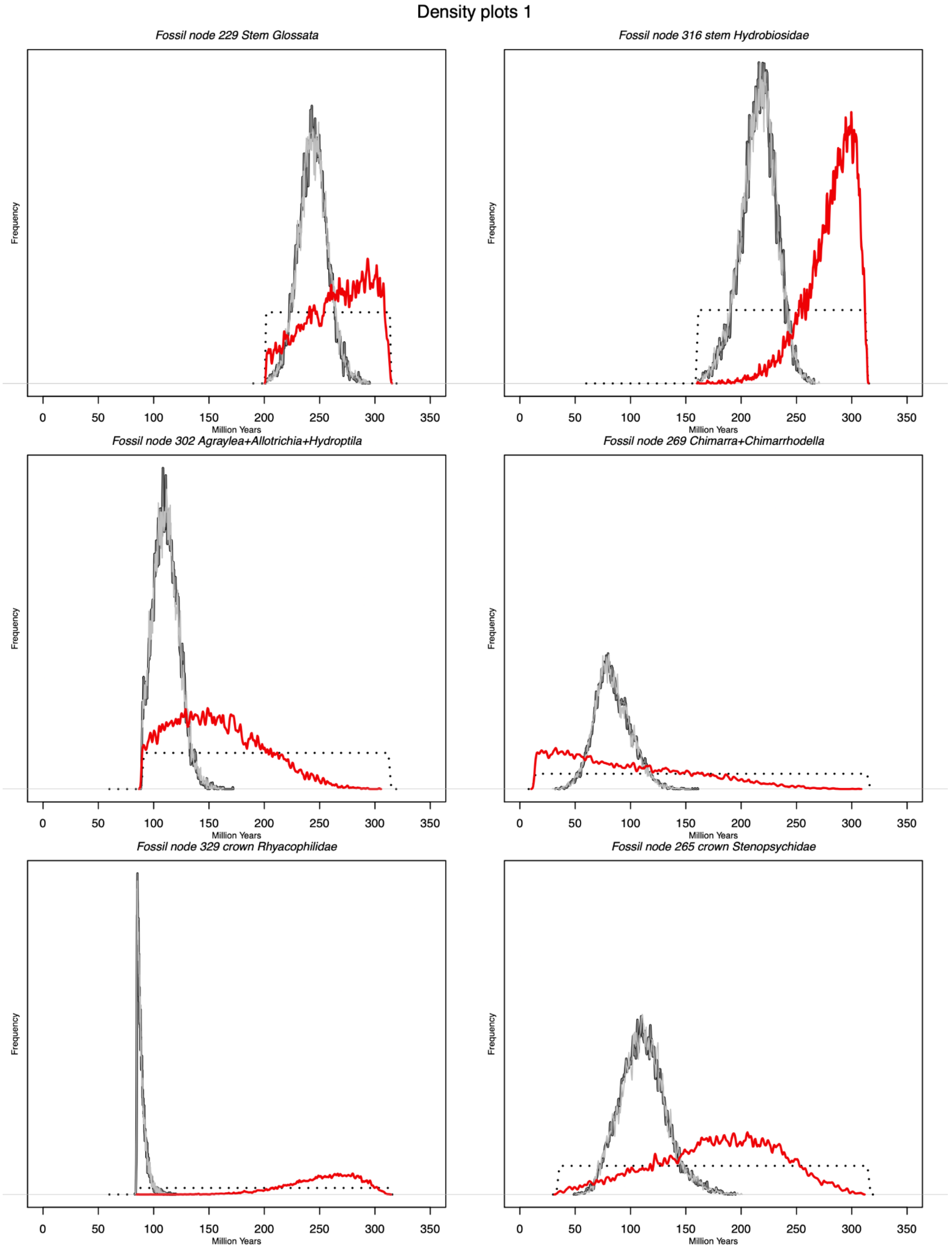

#### Density plots 2

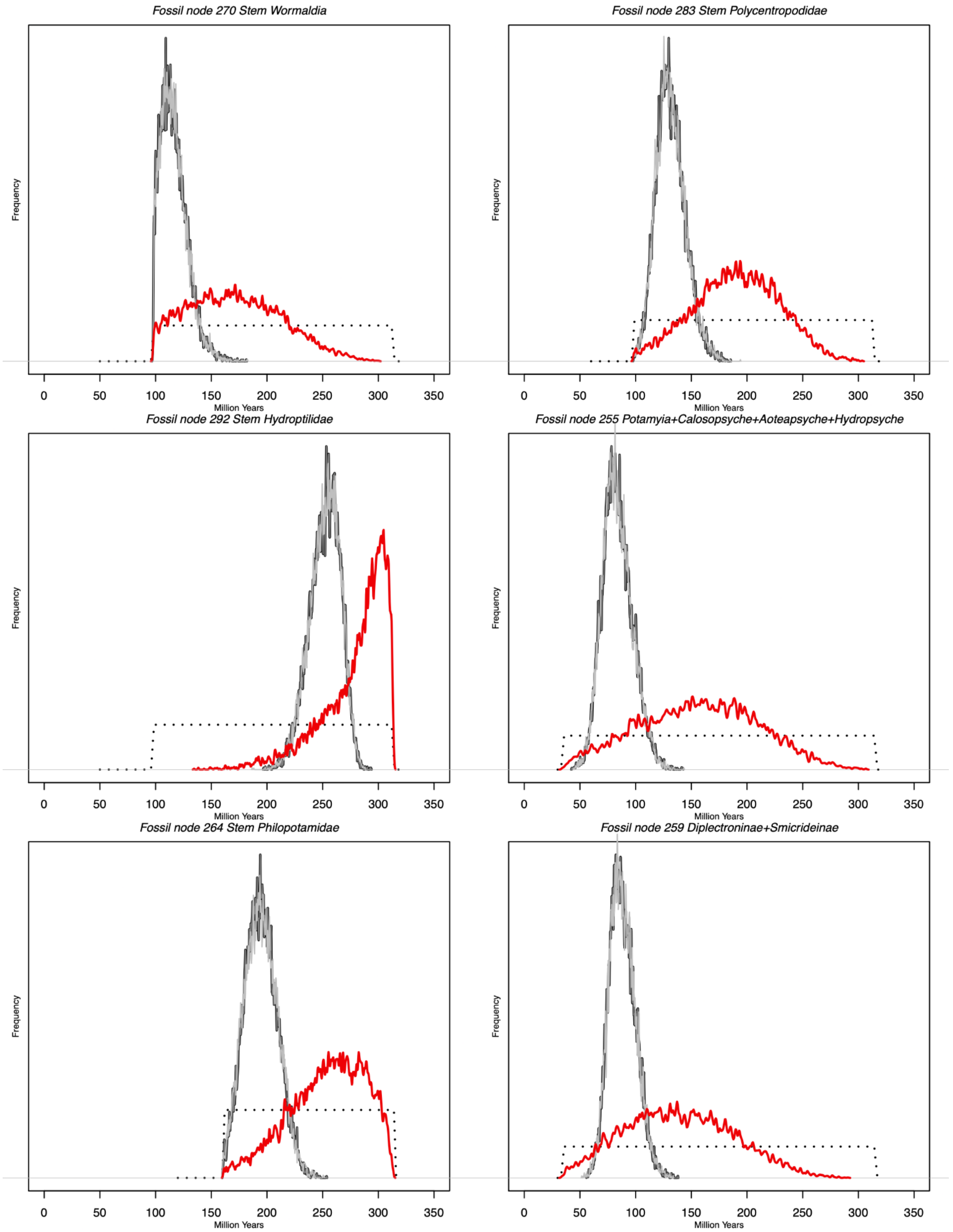

### Density plots 3

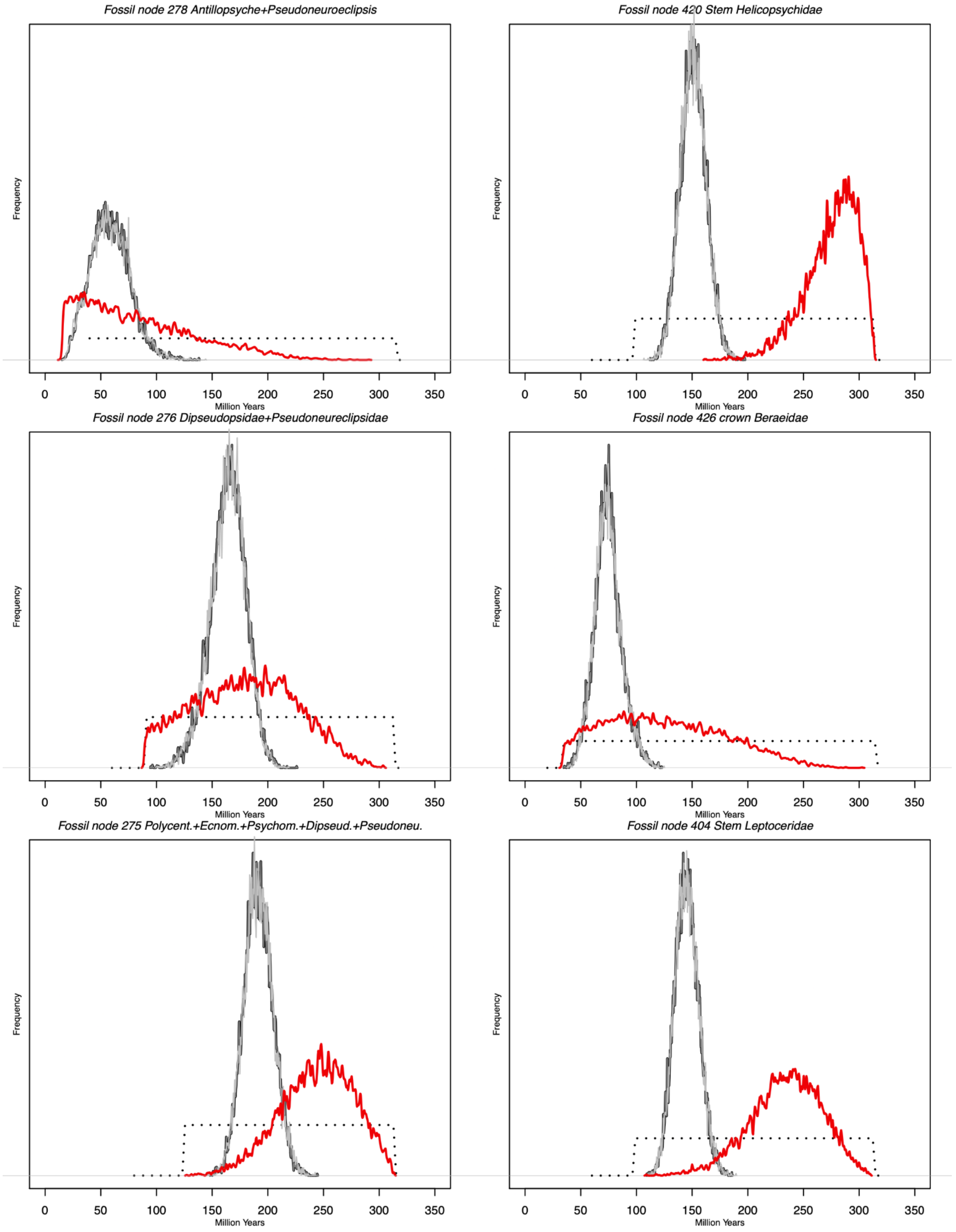

#### Density plots 4

*Fossil node 399 Stem Calamoceratidae*

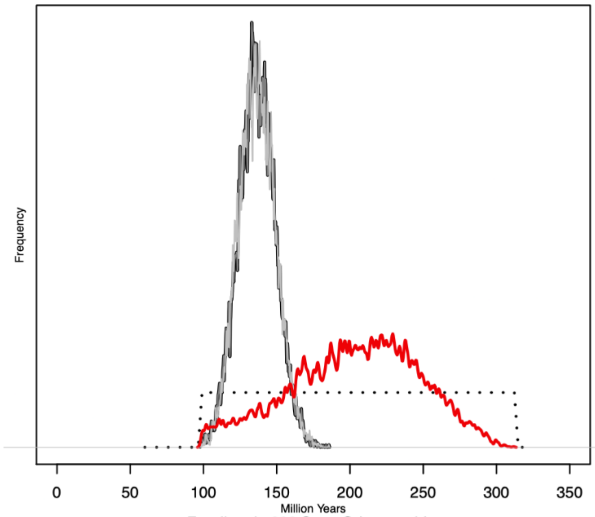


*Fossil node 409 Ceraclea+Brachysetodes+Athripsodes*

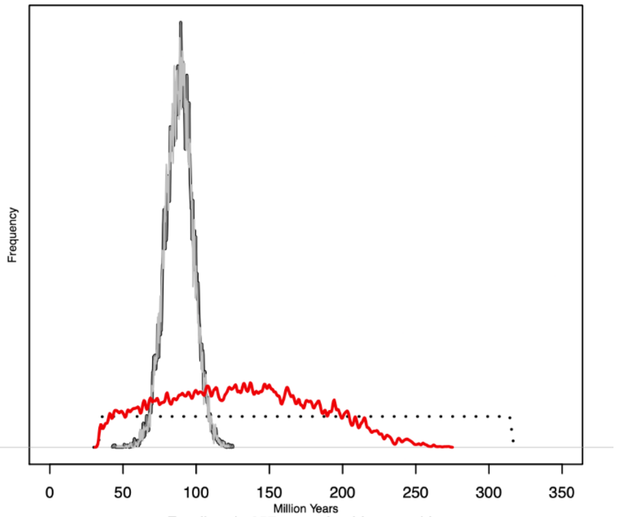


*Fossil node 392 Stem Odontoceridae*

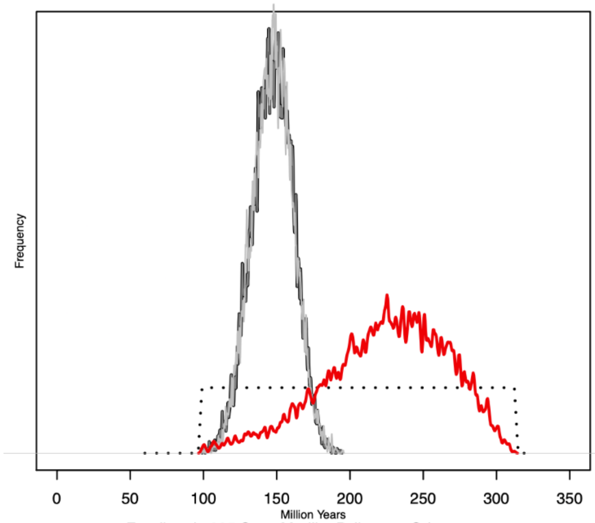


*Fossil node 357 crown Lepidostomatidae*

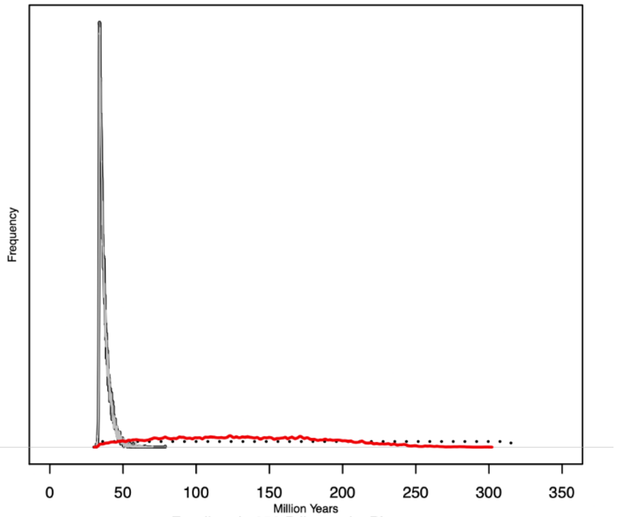


*Fossil node 395 Stem Marilia+Psilotreta+Odontocerum*

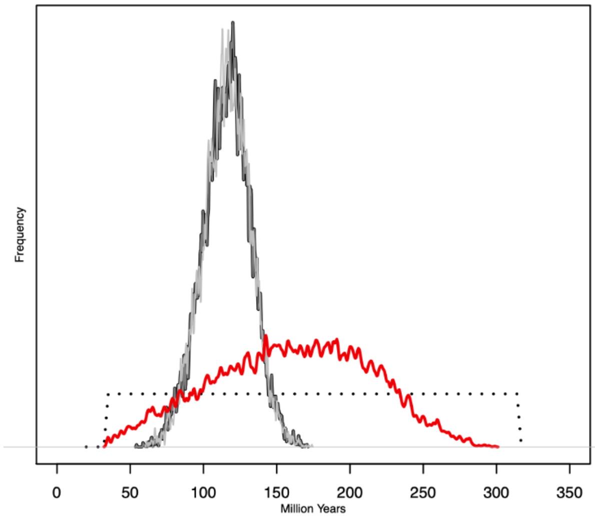


*Fossil node 353 Ptilostomis+Phryganea*

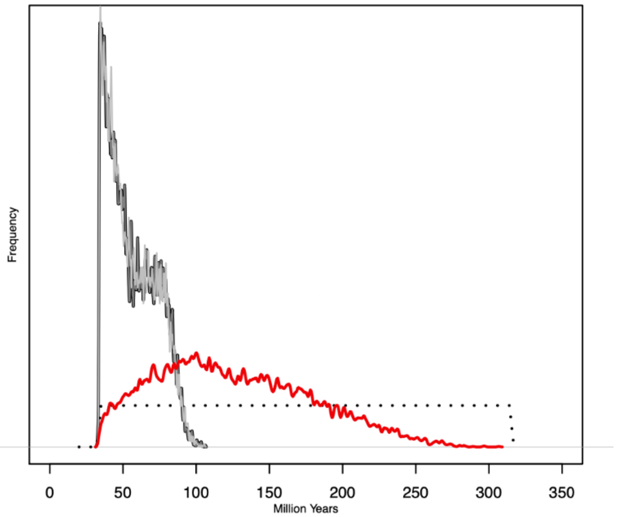

#### Density plots 5

*Fossil node 359 crown Brachycentridae*

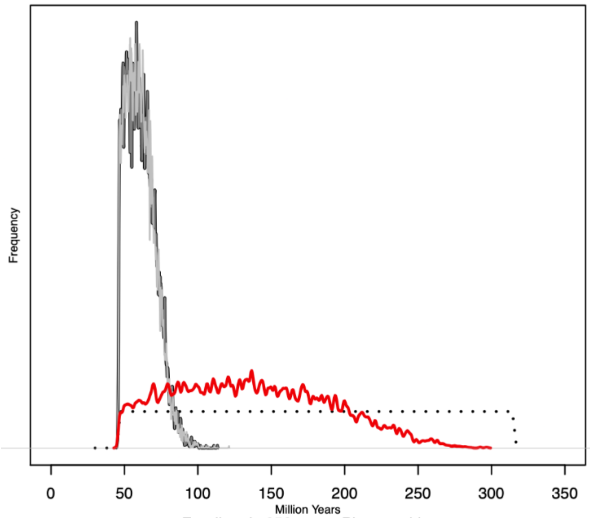


*Fossil node 419 Setodes+Mystacides*

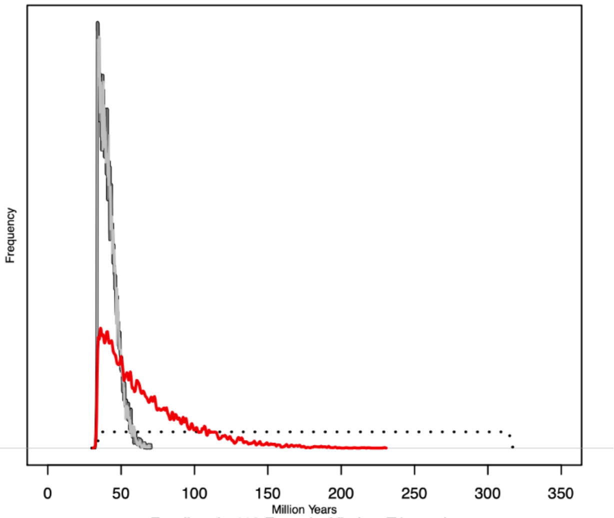


*Fossil node 352 crown Phryganeidae*

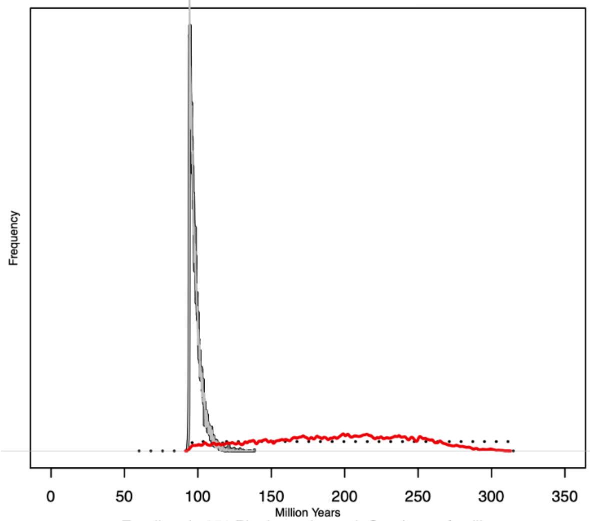


*Fossil node 416 Erotesis+Ylodes+Trianeodes*

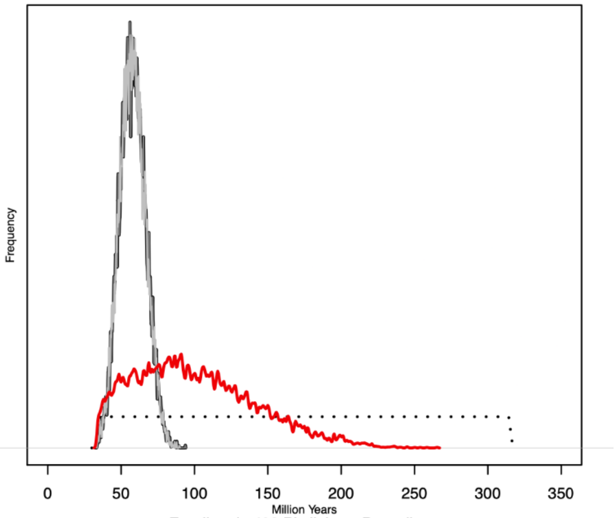


*Fossil node 351 Plenitentoria, excl. Gondwana families*

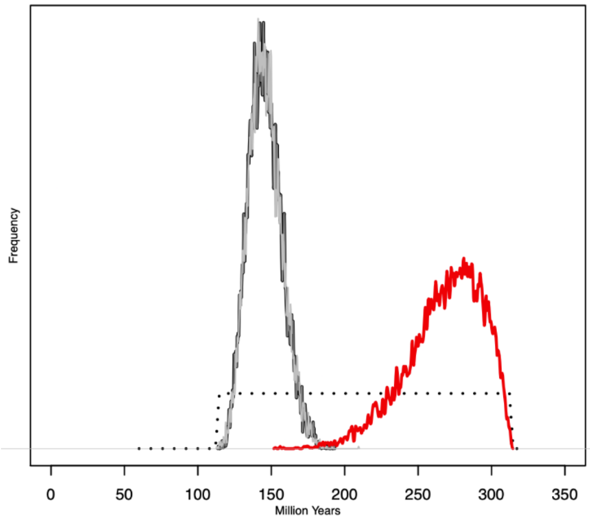


*Fossil node 403 Phylloicus+Banyallarga*

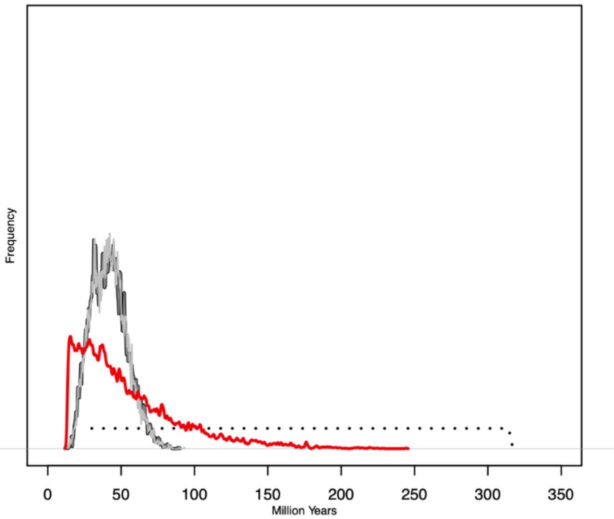

### Density plots 6

*Fossil node 288 Plectrocnemia+Polycentropus*

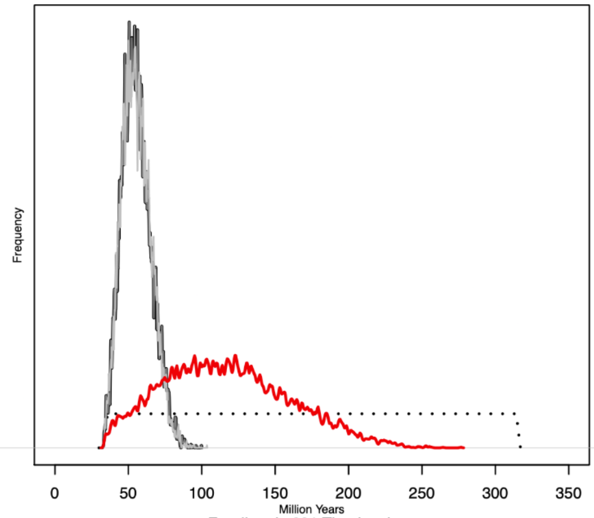


*Fossil node 281 Tinodes+Lype*

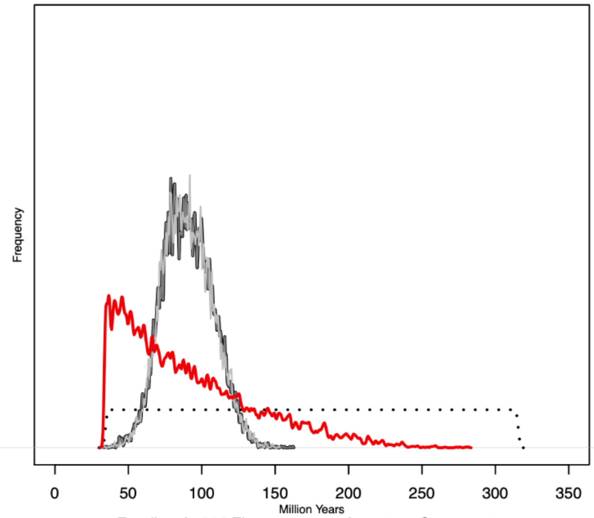


*Fossil node 314 Electragapetus+Agapetus+Synagapetus*

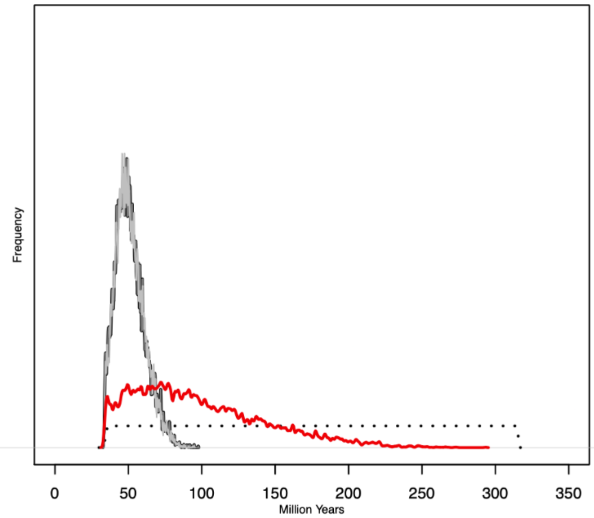

#### Fig. S6 Posterior mean ages plotted from separate MCMC dating runs with cauchy priors

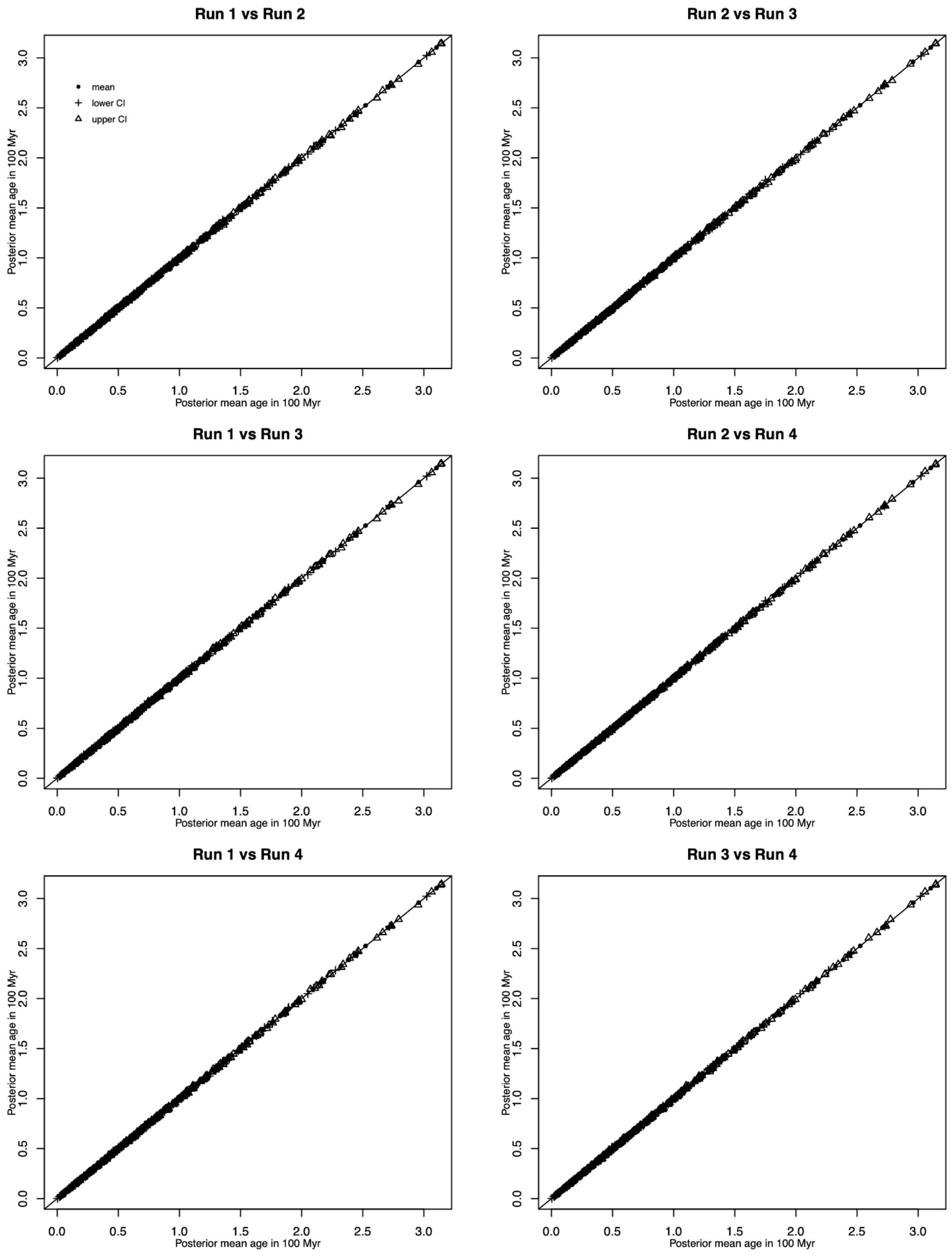

#### Fig. S6 Posterior mean ages plotted from separate MCMC dating runs with uniform priors

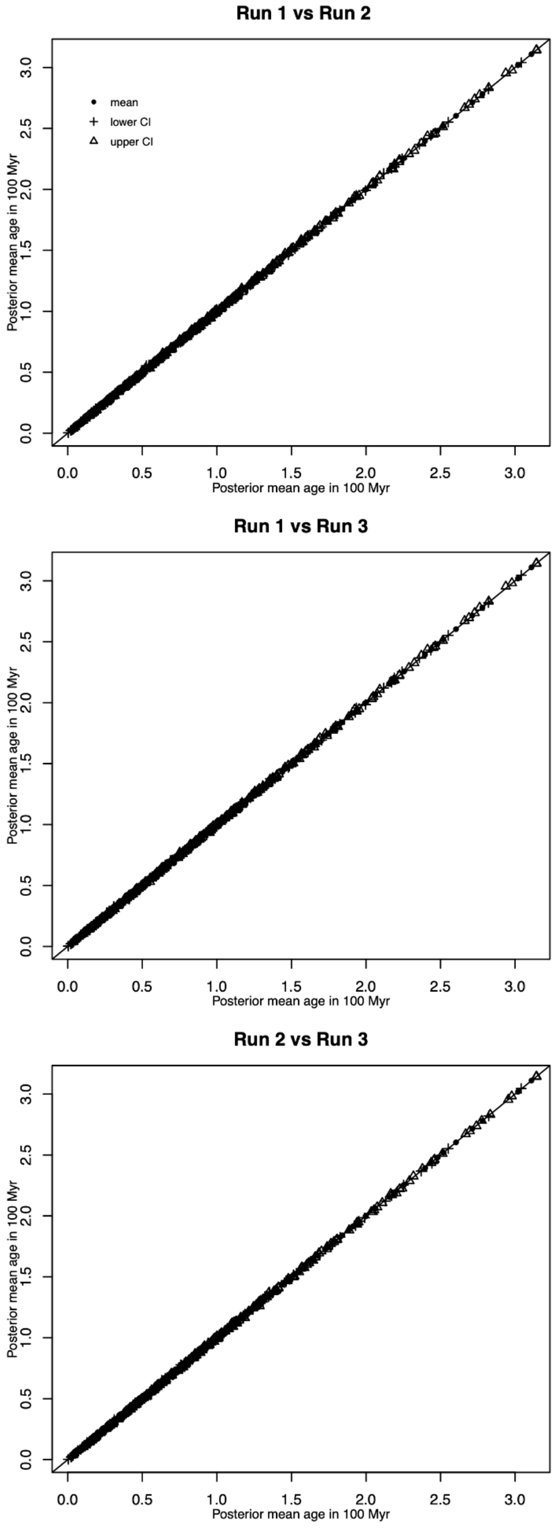
